# Structural basis of endogenous lipid recognition and G protein selectivity in GPR119

**DOI:** 10.64898/2026.08.27.747701

**Authors:** Donggyun Kim, Mohsen Ranjbar, Aleksey Raskovalov, Penghao Song, Jayden Salve, Vsevolod Katritch, Vadim Cherezov

**Affiliations:** Bridge Institute, Michelson Center for Convergent Bioscience, University of Southern California, Los Angeles, CA 90089, USA; Department of Chemistry, University of Southern California, Los Angeles, CA 90089, USA; Mork Family Department of Chemical Engineering and Material Science, University of Southern California, Los Angeles, CA 90089, USA; Department of Quantitative and Computational Biology, University of Southern California, Los Angeles, CA 90089, USA

**Keywords:** G protein-coupled receptor, GPR119, cryo-EM, Oleoylethanolamide (OEA), G protein selectivity, G_s_ protein, G_q_ protein

## Abstract

G protein-coupled receptors (GPCRs) often engage multiple intracellular transducers, yet the structural basis by which endogenous ligands influence G protein selectivity remains poorly understood. GPR119 is a lipid-activated receptor expressed in pancreatic β-cells and enteroendocrine L-cells, where it regulates glucose-dependent insulin and incretin secretion, making it a promising therapeutic target for metabolic disease. Here, we present cryo-electron microscopy structures of GPR119 bound to its endogenous lipid agonist oleoylethanolamide (OEA) in complex with G_s_ and G_q_ proteins. These structures reveal that OEA occupies a deeply buried orthosteric pocket but adopts distinct conformations in the two signaling states. Structural and functional analyses further identify an extended TM5 helix that stabilizes the receptor-G_s_ interface and acts as a key determinant of G-protein subtype selectivity. Together, these findings provide mechanistic insights into endogenous lipid-driven multi-transducer signaling and establish a structural framework for the development of pathway-selective GPR119 therapeutics.

## Introduction

Type 2 diabetes (T2D) is a chronic metabolic disorder characterized by insulin resistance and reduced insulin secretion from pancreatic β-cells, resulting in elevated blood glucose levels. The global prevalence of T2D is on the rise, largely driven by β-cell dysfunction associated with unhealthy lifestyle and obesity(*1*). GPR119 is an orphan class A G protein-coupled receptor (GPCR), often classified as a cannabinoid-like receptor together with GPR18 and GPR55, because these receptors respond to analogs of cannabinoid receptor ligands despite sharing minimal structural similarity with canonical cannabinoid receptors(*2*). GPR119 is expressed predominantly in pancreatic β-cells and intestinal enteroendocrine L-cells, playing critical roles in glucose homeostasis(*3*). Upon activation, GPR119 stimulates cyclic AMP (cAMP) production leading to glucose-dependent insulin secretion from pancreatic β cells and intestinal release of incretins, including glucagon-like peptide-1 (GLP-1) and glucose-dependent insulinotropic peptide (GIP)(*4–6*). Consequently, GPR119 has emerged as a promising therapeutic target for metabolic disorders, such as obesity and T2D(*7*).

Among several known endogenous agonists for GPR119, oleoylethanolamide (OEA) stands out as the most potent(*8*). OEA has positive effects in rats, including reducing food intake, reducing body weight(*9*), and modifying feeding behavior(*10*). Although GPR119 primarily couples to G_s_ protein, OEA has also been reported to induce calcium release and reduce intracellular cAMP levels through the activation of GPR119, indicating that OEA-mediated GPR119 activation is involved in multiple signaling pathways including those mediated by G_q_ and G_i_(*11*). The integrated signaling from multiple G proteins and other transducers is reported to play an important role in governing complex cell responses(*12*). In particular, both G_s_- and G_q_-mediated signaling have been strongly implicated in the regulation of peptide hormone secretion by lipid-sensing receptors. For example, the activation of both GPR119 and FFA1 (GPR40) through G_s_ and G_q_, respectively, acts in synergy to induce robust GLP-1 secretion(*13*). Although several structures of GPR119-G_s_ complexes bound to highly potent synthetic agonists have been reported(*14–16*), the molecular mechanisms underlying recognition and activation of GPR119 by endogenous ligands, as well as its coupling to G_q_ protein, remain poorly understood. Consequently, the structural basis for GPR119 coupling to multiple signaling transducers under physiological conditions remains elusive, largely because the determinants of G protein selectivity driven by endogenous ligand binding have not been established. To address these questions, we sought to define G protein coupling selectivity of GPR119 by endogenous lipid OEA.

Here, we determined two cryo-EM structures of OEA-bound GPR119 in complex with G_s_ and G_q_ proteins. These structures, along with molecular dynamics (MD) simulation and functional assays, revealed the molecular basis of GPR119 recognition of its endogenous ligand and coupling to distinct G protein subtypes, uncovering both shared and subtype-specific interaction networks at the receptor–G protein interface. Comparative structural analyses identified key residues that differentially contribute to G_s_ and G_q_ coupling, providing a mechanistic framework for understanding G protein selectivity driven by an endogenous lipid agonist. Together, our findings elucidate the structural determinants underlying multi-transducer coupling of GPR119 and offer mechanistic insights into how endogenous lipid ligands orchestrate diversified signaling. These results not only advance our understanding of GPR119-mediated signal transduction but also provide a structural foundation for the rational design of therapeutics targeting metabolic disorders such as T2D and obesity.

## Results

### 1. Structure determination of OEA-bound GPR119-G protein complexes

To facilitate the expression and purification of the human wild-type (WT) GPR119, we introduced a hemagglutinin signal peptide, FLAG tag, 10×His tag, and thermally stabilized bRIL(*17*) at the N-terminus, and a cleavable enhanced green fluorescent protein (eGFP) at the C-terminus of the canonical GPR119 gene sequence (UniProt Q13258). We also employed the Nanobit technology(*18*) to tether the C-terminus of GPR119 fused to LgBiT to the N-terminus of the G_β1_ subunit fused to HiBiT. The GPR119 construct was inserted into a pFastBac1 vector and codon-optimized for insect cell expression. The engineered GPR119 construct was co-expressed with the dominant-negative Gα_s_ (DNGα_s_)(*19*) or the engineered miniGα_q_(*20*) and G_β1γ2_ in *Spodoptera frugiperda* (*Sf9*) insect cells. Hereafter, G_s_ and G_q_ refer to the heterotrimeric G proteins consisting of DNGα_s_ or miniGα_q_, respectively, together with G_β1_ and G_γ2_, when describing the corresponding cryo-EM structures, unless otherwise specified. The GPR119-G_s_ and -G_q_ complexes were stabilized by adding OEA in the presence of apyrase, and the nanobody 35 (Nb35)(*21*) was further added to stabilize the GPR119-G_s_ complex. The structures of the endogenous lipid agonist OEA-bound GPR119-G_s_-Nb35 and GPR119-G_q_ complexes were determined by single-particle cryo-EM at nominal resolutions of 2.72 Å and 3.05 Å, respectively (Fig. 1A, B, Supplementary Fig. S1 and S2, and Supplementary Table S1). The high-resolution cryo-EM density maps enabled accurate model building for both OEA-bound GPR119-G_s_-Nb35 and GPR119-G_q_ complexes (Supplementary Fig. S3). The G_s_ complex contains residues 4 to 300 of GPR119, except for the intracellular loop 3 (ICL3; residues 212-218), and most residues of the G_s_ protein heterotrimer and Nb35, except for the flexible α-helical domain of the Gα_s_ protein. The G_q_ complex contains residues 4 to 302 of GPR119, except for ICL3 (residues 199-216), and most residues of the miniG_q_ heterotrimer. The endogenous agonist OEA was modeled into well-defined densities in the orthosteric GPR119 ligand-binding pocket in both structures (Fig. 1A, B). Both structures exhibit a canonical composition, in which OEA binds to the orthosteric pocket and G proteins bind to the intracellular side of GPR119, and both GPR119 structures share similar conformations (RMSD = 0.6 Å), except for the length of the transmembrane helix 5 (TM5) (Fig. 1A, B, and Supplementary Fig. S4).

**Figure 1.**
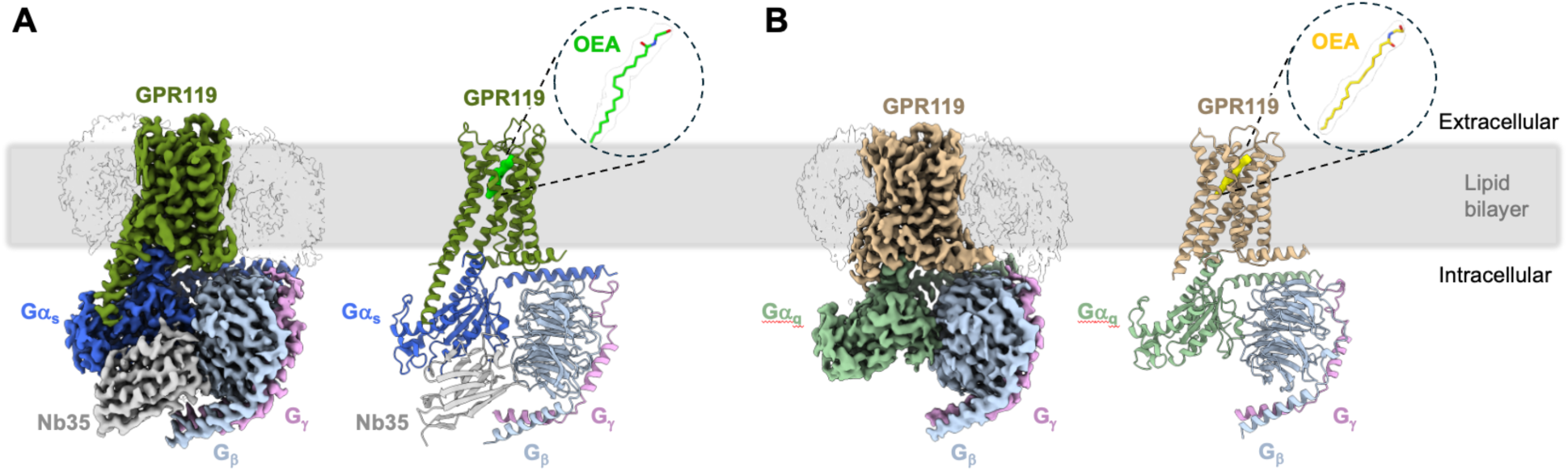
Cryo-EM structures of OEA-bound GPR119 coupled to G_s_ and G_q_. (**A**) Cryo-EM density map (contoured at a level 7) and atomic model of the OEA-bound GPR119-G_s_ complex, showing GPR119 (green), Gα_s_ (blue), G_β_ (light blue), G_γ_ (pink), and the stabilizing nanobody Nb35 (gray). (**B**) Cryo-EM density map (contoured at a level 9) and atomic model of the OEA-bound GPR119-G_q_ complex, with GPR119 (tan), Gα_q_ (light green), G_β_ (light blue), and G_γ_ (pink). In both complexes, OEA occupies the orthosteric binding pocket within the transmembrane domain of GPR119 and stabilizes an active receptor conformation compatible with coupling to distinct G protein subtypes. The gray slab indicates the membrane boundaries as defined by the Orientation of Proteins in Membranes (OPM) database.

To gain additional insights, we performed MD simulations of both complexes using the same constructs employed for cryo-EM structure determination, as well as corresponding models in which the engineered constructs were replaced with the wild-type (WT) sequences. All simulations showed stable trajectories with minimal deviations from the cryo-EM structures (Supplementary Fig. S5 and S6).

### 2. OEA binding pocket in G_s_ and G_q_ complexes

OEA is an endogenous N-acylethanolamine lipid, derived from oleic acid, that plays an important role in suppressing appetite and modulating feeding behaviors(*22, 23*). OEA stimulates incretin hormone secretion, contributing to glucose homeostasis(*24*), and has been recognized as the most potent endogenous agonist of GPR119(*6*).

The effect of OEA on G protein signaling was evaluated using a BRET-based ONE-GO G_s_ activation assay and an HTRF IP-ONE G_q_ signaling assay. OEA exhibits approximately a 3-fold stronger potency for G_s_ activation (EC₅₀ = 1.2 μM) than for G_q_ signaling (EC₅₀ = 3.7 μM) (Supplementary Fig. S7; Supplementary Tables S2 and S3), suggesting that OEA is a G_s_-preferential endogenous ligand, although these pharmacological characteristics were measured in different assay systems.

In both the G_s_- and G_q_-coupled cryo-EM structures, OEA binds to GPR119 within a deeply buried orthosteric pocket formed by transmembrane helices (TMs) 2, 3, 5, 6, and 7 as well as ECL2 (Fig. 2A–E). The ligand occupies an elongated, tunnel-like cavity that is largely occluded from the extracellular solvent, leaving only a narrow access channel to the pocket entrance. The binding cavity can be divided into two regions that accommodate the OEA head group and its hydrophobic acyl chain, respectively. The ethanolamide head group is located near the extracellular region, whereas the aliphatic tail extends deeply into the receptor core, forming extensive hydrophobic contacts along a curved tunnel. Notably, although the overall architecture of the pocket is highly similar between the two signaling states, OEA reaches approximately 2.1 Å deeper into the receptor core in the G_s_-coupled structure than in the G_q_-coupled structure (Fig. 2A, B). Consistent with this, comparison with previously reported GPR119-G_s_ structures bound to other ligands (*14, 15, 25*) shows that OEA penetrates the deepest into the receptor core (Supplementary Fig. S8).

**Figure 2.**
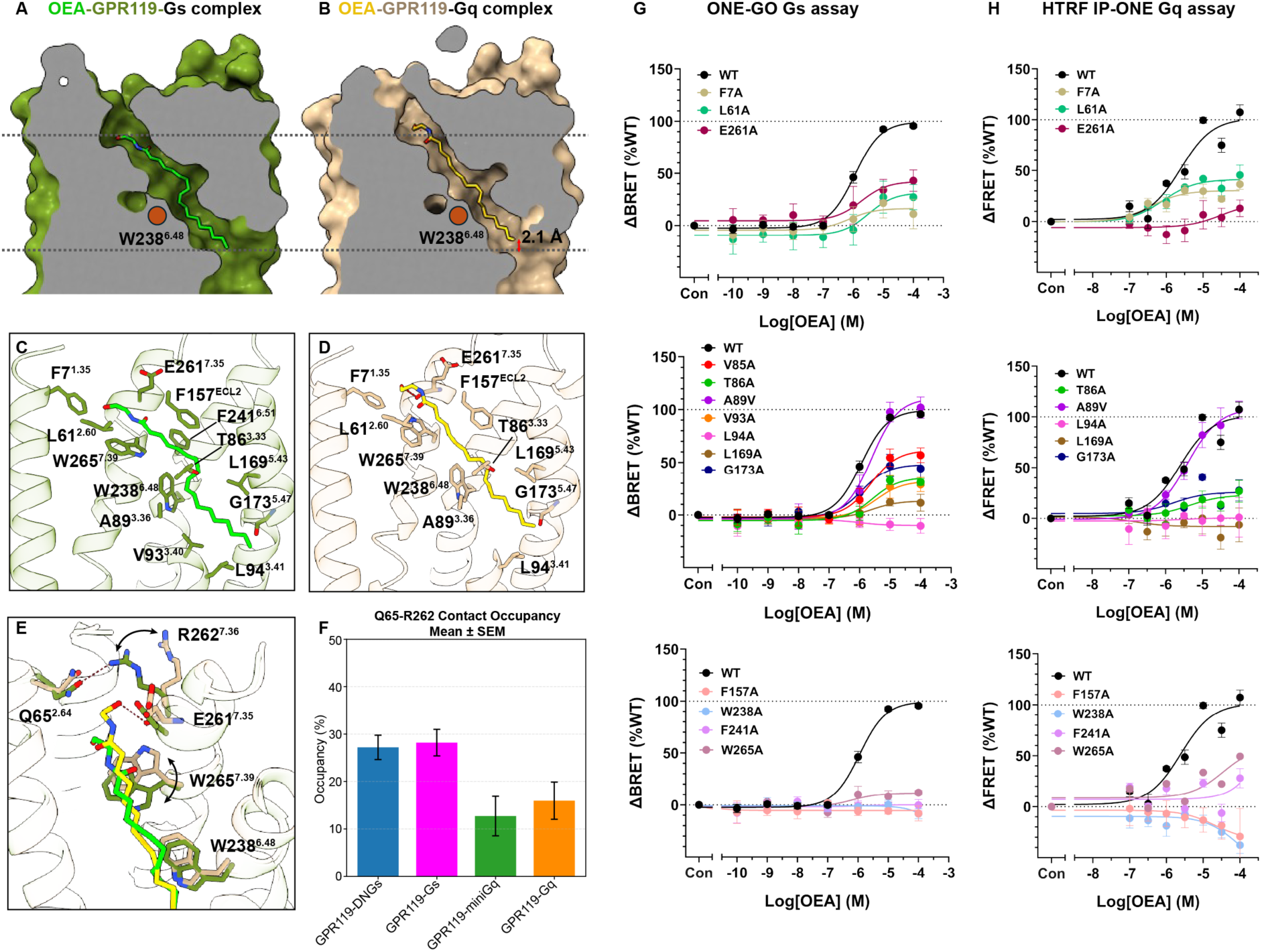
Architecture of the OEA binding pocket in GPR119. (**A**, **B**), Split surface representations of the OEA binding pocket in the G_s_- (green) and G_q_-coupled (wheat) GPR119 structures, showing a deeply buried, tunnel-like cavity accommodating the elongated OEA molecule. The position of the conserved toggle switch W238^6.48^ is indicated. The two black dotted lines denote the upper and lower boundaries of OEA in the G_s_-coupled structure and are used as reference planes to compare the ligand binding depth with that in the G_q_-coupled structure. (**C**, **D**) Detailed views of the OEA binding site in GPR119-G_s_ (C, green) and GPR119-G_q_ (D, wheat). OEA is shown as sticks and colored as lime or yellow, respectively, and the key residues are labeled. (**E**) Superimposed view of OEA binding sites showing distinct features between the OEA recognition residues in two complexes. (**F**) Comparison of the Q65^2.64^-R262^7.36^ contact occupancy (defined as the percentage of MD simulation frames where the closest heavy-atom distance between these two residues is ≤ 4 Å) among all structures. The bars show the average across all trajectories of each structure with error bars representing ±SEM. (**G**, **H**) Mutagenesis signaling assays for key residues in OEA binding sites. The ONE-GO G_s_ assay (G) and HTRF IP-ONE G_q_ assay (H) show concentration-dependent response curves induced by OEA at WT GPR119 with key residue mutants in G_s_ and G_q_ coupling, respectively. Data are normalized to WT maximal response and correspond to means ± SEM for three biologically independent experiments conducted in triplicates. The corresponding pEC_50_ and E_max_ values are shown in Supplementary Tables S2 and S3.

In the G_s_-coupled structure, the amide group of OEA is aligned parallel to the indole ring of W265^7.39^, allowing the amide–π interaction to stabilize the ethanolamide head group (Fig. 2C). The head group is further confined by hydrophobic interactions with F7^1.35^, L61^2.60^, and E261^7.35^ at the pocket entrance, which together shape the upper pocket of the binding site. In the G_q_-coupled structure, the ethanolamide head group of OEA is shifted ∼2.8 Å toward the extracellular side and TM7 relative to the G_s_-coupled structure and is also stabilized by F7^1.35^, L61^2.60^, and E261^7.35^ (Fig. 2D). Notably, in the G_q_-coupled complex, the hydroxyl group of OEA engages in a polar interaction with the backbone carbonyl of E261^7.35^, while W265^7.39^ adopts a flipped conformation to maintain its interaction with the ligand (Fig. 2E). Consistent with these structural observations, the alanine mutation of W265^7.39^ fully abrogates the activity of GPR119 in G_s_ signaling pathway and substantially impairs the G_q_ signaling activity (Fig. 2G, H). Moreover, alanine mutations of F7^1.35^, L61^2.60^, and E261^7.35^ markedly reduce the activity of GPR119 in both G_s_ and G_q_-mediated signaling pathways. Notably, E261^7.35^A exhibits a more pronounced effect on G_q_-mediated signaling compared to G_s_ (Fig. 2G, H), suggesting a preferential role of E261^7.35^ in OEA recognition within the G_q_ signaling pathway due to closer proximity of E261^7.35^ to the OEA headgroup in the GPR119-G_q_ structure (Fig. 2E).

The aliphatic acyl chain of OEA extends downward into a deeply buried, curved hydrophobic tunnel. Along this tunnel, OEA makes extensive van der Waals contacts with residues from ECL2 and TMs 3, 5, and 6, including F157^ECL2^, A89^3.36^, V93^3.40^, L94^3.41^, L169^5.43^, G173^5.47^, W238^6.48^, and F241^6.51^ (Fig. 2C, D). These residues collectively define the tunnel-like shape of the orthosteric pocket and tightly accommodate the hydrophobic tail of OEA. The deeper insertion of OEA in the pocket in the G_s_ complex relative to G_q_ is facilitated by the positioning of the C9-C10 double bond near the toggle switch residue W238^6.48^, enabling a strong π-π interaction. This interaction promotes a tighter fit into the hydrophobic pocket, with V93^3.40^ and L94^3.41^ potentially contributing to the enhanced ligand potency toward G_s_ pathway compared to G_q_. In contrast, in the G_q_ complex, the OEA double bond is shifted up by ∼2.0 Å and rotated compared to that of G_s_ complex, resulting in a weaker interaction with W238^6.48^ and a reduced penetration of the OEA acyl chain (Fig. 2C, D). Mutations of the hydrophobic residues V93^3.40^, L94^3.41^, L169^5.43^, and G173^5.47^ significantly reduce signaling activity in both pathways (Fig. 2G, H). Notably, the alanine substitution of L94^3.41^, which interacts with the terminal region of OEA, results in a complete loss of both G_s_- and G_q_-mediated activation, whereas the valine mutation of A89^3.36^ shows no marked effect on the activation of either G protein. In addition, the alanine mutants of the key aromatic residues F157^ECL2^, W238^6.48^, F241^6.51^, and W265^7.39^ almost completely abolish both G protein activation. These results indicate that aromatic residues play an important role in shaping the core pocket for ligand binding and make critical contributions to OEA recognition and subsequent G protein activation.

Detailed structural comparison of the OEA binding pockets between the two structures reveals that distinct ligand-binding modes lead to different conformations near the extracellular region and the upper part of TM7, particularly involving residues W265^7.39^ and R262^7.36^ (Fig. 2E). First, in the G_s_ complex, residue W265^7.39^ engages in an amide–π interaction with the ethanolamine carbonyl of OEA. In contrast, in the G_q_ complex, this residue adopts a flipped conformation, resulting in a shift of the OEA head group toward the extracellular side and TM7. The distinct conformations of W265^7.39^ remained stable throughout all MD simulations (Supplementary Fig. S9). Second, in the G_s_ complex, R262^7.36^ forms a hydrogen bond with Q65^2.64^, and despite lacking direct interaction with OEA, occludes the entrance of the extracellular pocket. In G_q_ complex, the hydroxyl head group of OEA protrudes out of the pocket disrupting the R262^7.36^ - Q65^2.64^ interaction and instead making a hydrogen bond with the backbone carbonyl of E261^7.35^ residue. This leads to an outward rotation of R262^7.36^, resulting in a more open extracellular pocket entrance compared with the G_s_ complex (Fig. 2E). MD simulations supported more persistent interactions between R262^7.36^ and Q65^2.64^ in the G_s_ complex (Fig. 2F). Additionally, analysis of the frequency of direct contacts between the ligand and receptor residues across all MD simulations revealed distinct interaction patterns involving L61^2.60^ and F7^1.35^ residues in the G_s_ and G_q_ complexes (Supplementary Fig. S10).

### 3. Comparison of G protein-binding interface

While GPR119 primarily activates the G_s_ signaling pathway, it has also been reported to couple to multiple downstream effectors upon activation(*11*). To investigate the mechanism underlying G_s_- and G_q_-mediated signaling by GPR119, we compared our high-resolution cryo-EM structures. Based on receptor alignment, G_s_- and G_q_-coupled GPR119 structures exhibit nearly identical conformations with a Cα root mean square deviation (RMSD_Cα_) of 0.6 Å within the 7TM bundle (Supplementary Fig. S4). The most pronounced difference is observed in TM5, which is extended by approximately 3 helical turns in the G_s_-coupled GPR119 structure compared to the G_q_-coupled one (Fig. 3A). This extension enables additional interactions with the Ras domain of Gα_s_, while the corresponding part of Gα_q_ does not interact with the receptor, as described in detail below.

**Figure 3.**
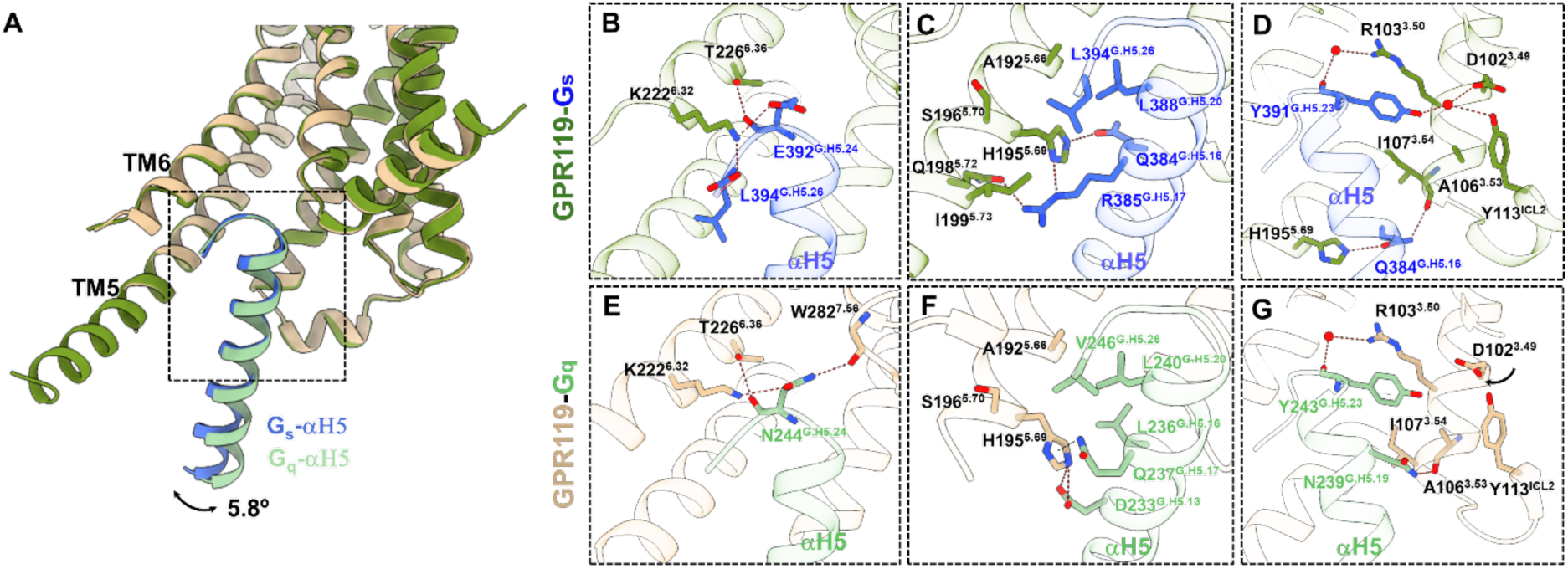
Distinct interactions of the Gα_s_ and Gα_q_ α5 helices with GPR119. (**A**) Overall superposition of GPR119 in complex with Gα_s_ (blue) and Gα_q_ (green), highlighting the relative orientation of the Gα α5 helix. The α5 helices closely overlap at their tips but adopt distinct tilt angles, differing by ∼5.8°. (**B**-**D**) Close-up views of the interface between the Gα_s_ α5 helix and the GPR119 transmembrane core, showing key polar interactions. (**E**-**G**) Corresponding close-up views of the Gα_q_ α5 helix–receptor interface.

At the G protein-binding interface in both structures, the C-terminal α5 helix of the Gα subunit inserts into an intracellular cavity formed by TM3, 5, 6, and 7 of GPR119. A 5.8° difference in the tilt angle of the α5-helix is observed between the Gα_s_ and Gα_q_ proteins (Fig. 3A). MD simulations suggest that the α5-helix exhibits greater conformational dynamics in the G_q_ complex, as reflected by the broader distribution of its tilt angle (Supplementary Fig. S11). The overall interaction patterns between the α5-helix and GPR119 residues observed in the MD simulations are similar in both complexes (Supplementary Fig. S12).

Of the 12 and 9 GPR119 residues that form polar interactions with the α5-helix of Gα_s_ and Gα_q_, respectively, 8 are shared between the two interfaces (Fig. 3B−G). The TM6 residues K222^6.32^ and T226^6.36^ of GPR119 form polar interactions with E392^G.H5.24^ and C-terminal carboxyl of L394^G.H5.26^ (superscript indicates the common Gα numbering (CGN) system(*26*)) in Gα_s_ and with N244^G.H5.24^ in Gα_q_. In addition, the backbone carbonyl of W282^7.56^ in Gα_q_ engages in a hydrogen bond with N244^H5.24^ (Fig. 3B, E). The TM5 residues H195^5.65^ and Q198^5.72^ form polar interactions with Q384^G.H5.16^ and R385^G.H5.17^ of Gα_s_, while A192^5.66^, S196^5.70^, and I199^5.73^ on TM5 form hydrophobic interactions with Gα_s_ residues L394H^G.5.26^, L388^G.H5.20^, R385^G.H5.17^ (Fig. 3C). Meanwhile, the residue H195^5.65^ makes a polar interaction with D223^G.H5.13^ and an NH-π interaction with the side-chain NH group of Q237^G.H5.17^ in Gα_q_, whereas Q198^5.72^ and I199^5.73^ do not interact with Gα_q_ (Fig. 3F). The conserved residue R103^3.50^ within the DRY motif, a key activation switch, engages in π-stacking interactions with the sidechain of Y391^G.H5.23^ and forms a water-mediated hydrogen bond with the backbone carbonyl of Y391^G.H5.23^ in both structures. In addition, the backbone carbonyls of I107^3.54^ and A106^3.53^ form polar interactions with Q384^G.H5.16^ of Gα_s_ and N239^G.H5.19^ of Gα_q_, respectively (Fig. 3D, G). Notably, the sidechain of Y391^G.H5.23^ in Gα_s_ participates in a water-mediated polar interaction network involving the conserved DRY-motif residue D102^3.49^ and Y113^ICL2^. In contrast, this polar network is absent in the G_q_ complex, due to the lack of the bridging water molecule, accompanied by a reorientation of the D102^3.49^ side chain (Fig. 3D, G).

The αN helix and β2–β3 loop of Gα_s_ are positioned 3.5−6.0 Å closer to the intracellular cavity of GPR119 than the corresponding regions of Gα_q_, resulting in distinct interaction modes between the intracellular loops of GPR119 and the G_s_ and G_q_ heterotrimers (Fig. 4A). The ICL2 residue K115^ICL2^ in both G_s_- and G_q_-coupled GPR119 stabilizes the G protein interface by a polar interaction with the backbone carbonyl of K216^G.S3.00^ (Fig. 4B, D). The K115^ICL2^A mutation significantly reduces the activation of both G proteins, suggesting this interaction is crucial for the recruitment of both G_s_ and G_q_ proteins (Fig. 4E, F). In addition, the sidechain of F111^ICL2^ is buried into a hydrophobic pocket formed by I383^G.H5.15^ R380^G.H5.12^, F376^G.H5.08^, V217^G.S3.01^, and H41^G.S1.02^ in Gα_s_ (Fig. 4B). Although the corresponding residues of the Gα_q_ α5 helix form hydrophobic interactions with F111^ICL2^, no interactions are observed with β1 and β3 of Gα_q_, owing to the less engaged position of G_q_ at GPR119 (Fig. 4D).

**Figure 4.**
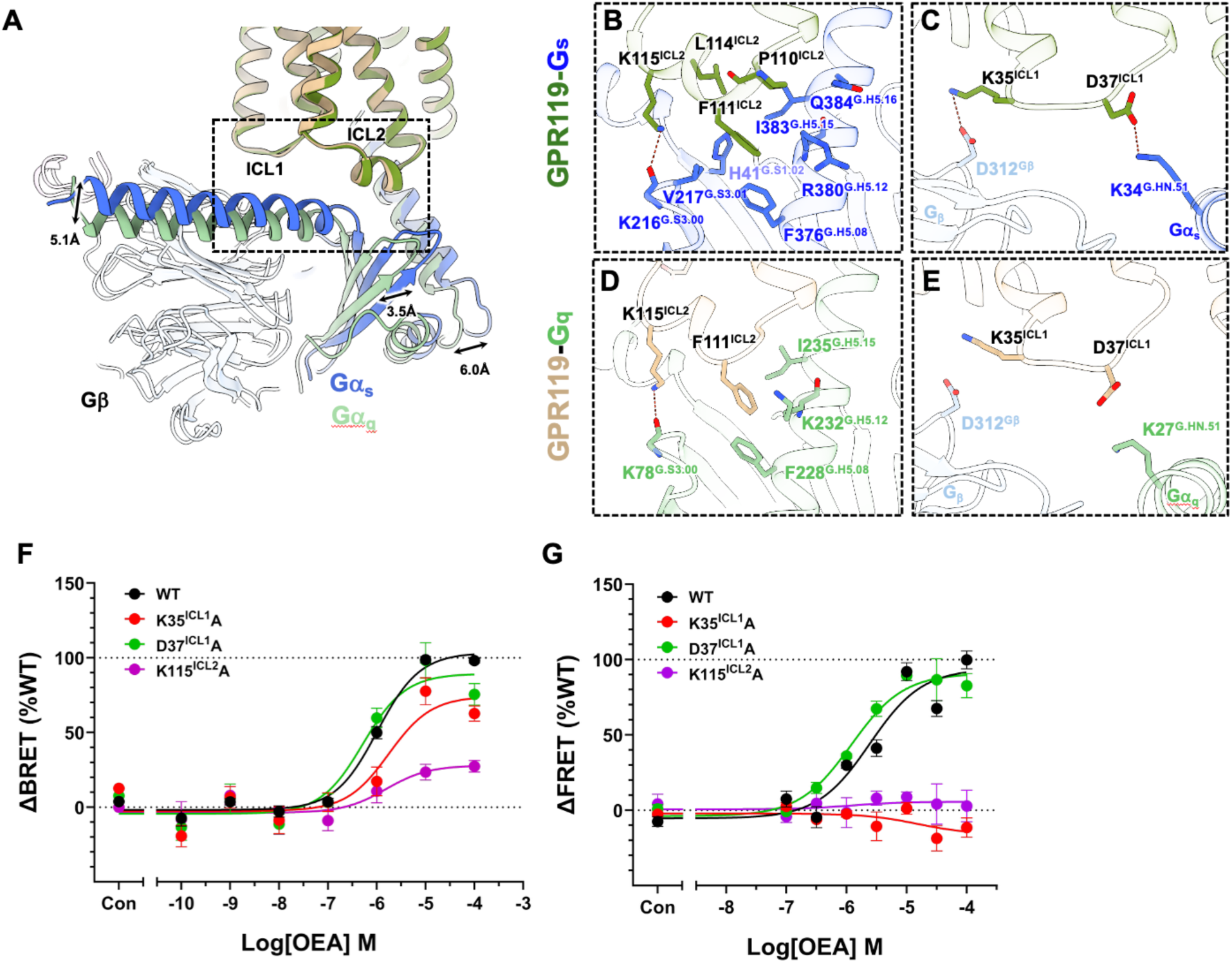
Comparison of G protein binding interface with intracellular loops of GPR119. (**A**) Structural superposition of the GPR119-Gα_s_ and GPR119-Gα_q_ complexes, highlighting differences in the intracellular loop regions (ICL1 and ICL2) and overall G protein–coupling interface. (**B**, **C**) Enlarged views of the interface between Gα_s_ and ICL2 (B) and ICL1 (C). (**D**, **E**) Enlarged views of the interface between Gα_q_ and ICL2 (D) and ICL1 (E). The red-dotted lines indicate polar interaction. (**F**, **G**) Concentration-dependent response curves of WT GPR119 and its G protein-interface mutants in response to OEA using the ONE-GO G_s_ assay (F) and the HTRF IP-ONE G_q_ assay (G). Data were normalized to WT maximal response and correspond to means ± SEM for three biologically independent experiments conducted in triplicates. The pEC_50_ and E_max_ values are provided in Supplementary Tables S2 and S3.

The two ICL1 residues K35^ICL1^ and D37^ICL1^ make salt bridges with D312 of Gβ subunit and K34^G.HN.51^ of Gα_s_ in GPR119-G_s_ complex, respectively (Fig. 4C), whereas in the GPR119-G_q_ complex, both of these interactions have longer distances (Fig. 4E). The D37^ICL1^A mutant shows no significant effect on G_q_ activation and a modest reduction in G_s_ signaling efficacy, indicating that this residue preferentially contributes to G_s_ coupling, consistent with our structural analysis, although its impact is not substantial (Fig. 4F, G). Notably, the K35^ICL1^A mutant completely abolishes G_q_ signaling and substantially reduces G_s_ signaling (Fig. 4F, G). This observation may be explained by the comparatively sparse interaction network at the G_q_ interface, rendering G_q_ coupling more sensitive to single-residue substitutions than G_s_ coupling (Supplementary Fig. S13).

### 5. Contribution of TM5 to G protein selectivity

Next, we performed a detailed analysis of the interactions between TM5 of GPR119 and each G protein. The most striking difference between G_s_- and G_q_-coupled GPR119 complexes is the presence of an extended TM5 in the G_s_ complex (Fig. 5A). These three additional helical turns in TM5 (residues 199–210) are well resolved with strong map density exclusively in the G_s_ complex, whereas the corresponding region in the G_q_ complex is completely disordered (Fig. 5A−C). In the G_s_ complex, the extended TM5 forms extensive interactions with the Ras-like domain of Gα_s_. Notably, two polar interactions, Q198^5.72^-R385^G.H5.17^ and K201^5.75^-D323^G.hgh.4.1^, are observed at the interface with Gα_s_ (Fig. 5B). Although the residue Q198^5.72^ is present in both complexes, the corresponding residue to Gα_s_ R385^G.H5.17^ is substituted with Q237^G.H5.17^ in Gα_q_, resulting in the absence of the analogous polar interaction.

**Figure 5.**
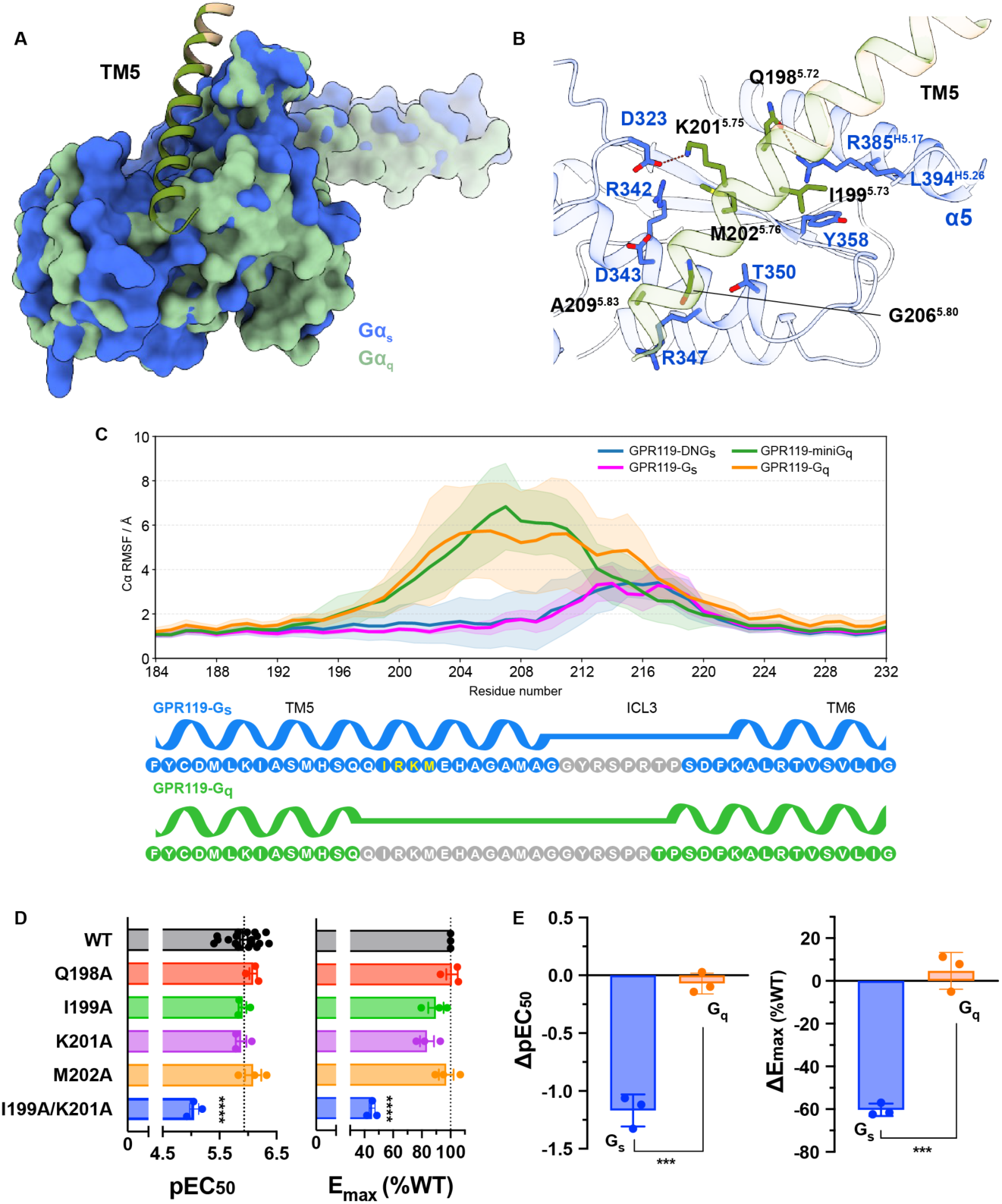
Structural and functional analysis of TM5 contribution to G protein selectivity. (**A)** Superposition of the TM5–Gα interface in the GPR119-G_s_ and GPR119-G_q_ complexes. (**B**) Detailed interactions between GPR119 TM5 and Gα_s_. Residues involved in the interactions are shown as sticks. Polar interactions are depicted as red dotted lines. **(C)** RMSF values for the TM5−ICL3−TM6 region obtained from MD simulations support longer stable TM5 in GPR119-G_s_ complex compared to GPR119-G_q_. The residues with well-defined density modeled in cryo-EM structures along with their secondary structure are highlighted in blue (GPR119-G_s_) and green (GPR119-G_q_), while the disordered residues are in grey. The key residues in TM5 region are colored yellow. (**D**) OEA-induced Gα_s_ signaling for WT GPR119 and TM5 mutants, measured by ONE-GO assay. The obtained potency (pEC_50_) and efficacy (E_max_) values for mutants were compared with those of WT by one-way ANOVA with Dunnett’s multiple comparisons test (p < 0.0001****). **(E)** Distinct effects of the I199^5.73^A/K201^5.75^A double mutant on two G protein signaling pathways. The double mutant selectively impairs the G_s_-mediated signaling while preserving the G_q_-mediated response. Deviations in potency and efficacy from those of the WT receptor are presented as mean ± SEM from three independent experiments. Significant difference was determined by two tail t-test (p < 0.001***). Concentration-response curves related to panels (D) and (E) are shown in Supplementary Fig. S14, with pEC_50_ and E_max_ values provided in Supplementary Tables S2 and S3.

In addition to these polar interactions, I199^5.73^ and M202^5.76^ of GPR119 exhibit hydrophobic interactions with R385^G.H5.17^**/**L394^G.H5.26^/Y358^G.h4s6.20^ and R342^G.H4.12^/L346^G.H4.16^, respectively (Fig. 5B). The C-terminal end of TM5, comprising G206^5.80^ and A209^5.83^ is further stabilized by the interface with D343^G.H4.13^, L346^G.H4.16^, R347^G.H4.17^, and T350^G.h4s6.03^ (Fig. 5B). Although single mutants Q198^5.72^A, I199^5.73^A, K201^5.75^A, and M202^5.76^A have only a minimal effect on OEA-induced G_s_ signaling, the combined I199^5.73^A/K201^5.75^A mutant results in an 8-fold reduction in potency and a 2.5-fold decrease in efficacy (Fig. 5D). To investigate the contribution of these residues to G_q_ activation, the I199^5.73^A/K201^5.75^A double mutant was further evaluated in a G_q_ signaling assay. Strikingly, in contrast to its effect on G_s_ signaling, alanine substitutions at these two positions did not alter G_q_ activity (Fig. 5E), suggesting that TM5 plays a crucial role in G protein subtype selectivity, with I199^5.73^ and K201^5.75^ serving as key determinants of this selectivity.

### 6. Structural comparison with other GPCRs

To explore whether the structural features underlying G_s_ versus G_q_ coupling observed in GPR119 are conserved among class A GPCRs, we compared GPR119 with other receptors whose structures have been determined in complex with both G_s_ and G_q_ proteins. These include histamine-bound H2R-G_s_ and -G_q_ complexes (PDB entries: 8YN3 and 8YN4)(*27*), ZH8651-bound mTAAR1-G_s_ and -G_q_ complexes (PDB entries: 8WC4 and 8WC9)(*28*), DHA-bound FFAR4-G_s_ and -G_q_ complexes (PDB entries: 8H4I and 8H4L)(*29*), and CCK8-bound CCK1R-G_s_ and -G_q_ complexes (PDB entries: 7EZK and 7EZM)(*30*) (Supplementary Fig. S15A−E).

Although these receptors are capable of engaging multiple transducers, H2R and TAAR1 primarily couple to G_s_ proteins, whereas FFAR4 and CCK1R predominantly couple to G_q_ proteins. Notably, while GPR119 displays a marked shift in ligand binding mode between the G_s_ and G_q_ coupled complexes, these other receptors show no significant changes in ligand binding between the two different G protein complexes (Supplementary Fig. S15F−J). Consistent with our observations, the H2R-G_s_ complex exhibits an intact TM5–ICL3–TM6 region compared with the G_q_-coupled structures, and mTAAR1-G_s_ also shows an extended TM5 relative to the G_q_ structure. In addition, the α5 helix (αH5) and N-terminal α-helix (αN) of Gα_s_ are tilted more toward ICL2 than those of Gα_q_, potentially enabling stronger receptor-G_s_ interactions (Supplementary Fig. S15K−M). In contrast to these G_s_-preferential receptors, the G_q_-preferential receptors, FFAR4 and CCK1R, exhibit the opposite structural trend, with a more intact and well-defined TM5–ICL3– TM6 region observed in the G_q_-coupled structures than in their G_s_-coupled counterparts (Supplementary Fig. S15N, O). Sequence alignment analysis indicates that the specific residues responsible for G protein selectivity are not conserved across these receptors (Supplementary Fig. S16), suggesting that G protein subtype selectivity is achieved through receptor-specific structural determinants. Nevertheless, the conformation of the TM5–ICL3–TM6 region appears to play a crucial role in differential G protein coupling across Class A GPCRs. Taken together, these results suggest that the structural principle underlying G protein selectivity identified in this study may be broadly conserved across class A GPCRs, whereas the precise regulation of G protein subtype coupling is likely achieved through receptor-specific structural features.

## Discussion

This study provides structural and mechanistic insights into how an endogenous lipid agonist oleoylethanolamide (OEA) activates GPR119 and promotes coupling to distinct G protein subtypes. By determining cryo-EM structures of OEA-bound GPR119 in complex with G_s_ and G_q_ proteins, we reveal the molecular basis of endogenous ligand recognition and identify structural determinants underlying G protein coupling selectivity. While G_s_-coupled structures in complex with synthetic ligands and the endogenous lipid lysophosphatidylcholine (LPC) have been reported, this study presents the first cryo-EM structure of GPR119 bound to its most potent endogenous agonist, OEA, in complex with G_s_ protein, as well as the first structure of GPR119 in complex with G_q_ protein.

OEA binds within a deeply buried orthosteric pocket formed by TMs 2, 3, 5, 6, and 7 together with ECL2, occupying an elongated hydrophobic tunnel in the receptor core. Despite the overall similarity of the binding pocket, OEA adopts distinct conformations in the G_s_- and G_q_-coupled structures. In the G_s_ complex, the ligand penetrates deeper into the receptor core and engages in a carbonyl–π interaction with W265^7.39^, positioning its C9–C10 double bond near the conserved toggle switch residue W238^6.48^. In contrast, in the G_q_ complex, the W265^7.39^ sidechain is flipped resulting in a shift of the ethanolamide head group toward the extracellular region. This observation suggests that subtle differences in ligand positioning can propagate structural changes to the intracellular signaling interface. Consistently, mutagenesis demonstrates that aromatic and key hydrophobic residues shaping the pocket are critical for receptor activation.

A key structural distinction between the two signaling states is the conformation of TM5 and its contribution to the receptor-G protein interface. In the G_s_ complex, TM5 adopts an extended helical conformation that forms extensive interactions with the Ras-like domain of Gαs, whereas the corresponding region is disordered in the G_q_ complex. Residues within this extended TM5 region, particularly I199^5.73^ and K201^5.75^, substantially contribute to G_s_ activation but have minimal effect on G_q_ signaling, indicating that TM5 acts as a structural determinant of G protein subtype selectivity in GPR119.

Beyond the distinct TM5-mediated interactions, both complexes share conserved receptor-G protein interface features, including insertion of the Gα C-terminal α5 helix into the intracellular cavity and stabilization by the DRY motif. Nevertheless, Gα_s_ engages more extensively with intracellular loops than Gα_q_, which may contribute to preferential G_s_ coupling. Comparative analysis with other class A GPCRs reveals that the TM5–ICL3–TM6 conformation represents a recurring structural feature that influences G-protein selectivity.

Understanding how GPR119 differentially engages G_s_ and G_q_ signaling has implications for therapeutic development targeting metabolic disorders. Activation of G_s_ in pancreatic β-cells and enteroendocrine L-cells enhances cAMP production, promoting glucose-dependent insulin and incretin secretion which are beneficial for glycemic control(*31, 32*), whereas G_q_ signaling modulates intracellular calcium, membrane excitability, and hormone release, potentially contributing complementary or context-dependent metabolic effects(*33, 34*). Increasing evidence suggests that GPCR ligands capable of selectively modulating specific pathways can improve therapeutic efficacy while minimizing adverse effects (*35*). Furthermore, complementary activation of G_s_ and G_q_, as observed in studies with FFA1 and GPR119 agonists or with GLP-1 signaling in β-cells, may enhance metabolic outcomes (*13, 36, 37*), highlighting that both biased signaling and synergistic pathway engagement may provide opportunities for therapeutic development.

In conclusion, our structures define the molecular basis of endogenous lipid recognition and reveal how GPR119 engages distinct G protein subtypes. By capturing OEA-bound GPR119 in complex with both G_s_ and G_q_ proteins, this work provides structural insights into the mechanisms underlying G-protein coupling selectivity. These findings establish a framework for understanding multi-transducer signaling in GPR119 and may guide the design of pathway-selective modulators for metabolic diseases including T2D and obesity.

## Acknowledgments

We thank Jeffrey Velasques, Kelly Villers, and Delainey Landaker for assistance with baculovirus and mammalian cell cultures, Htet Khant for cryo-EM user support, and Katya Kadyshevskaya for help with illustrations. This work was supported by the National Institutes of Health grants R35GM127086 (V.C.) and R35GM153437 (V.K.). Cryo-EM data were collected at the Center of Excellence in NanoImaging (CNI) at the University of Southern California. We acknowledge the Center for Advanced Research Computing (CARC) of the University of Southern California for providing computational resources for Cryo-EM data processing and MD simulations.

## Author contributions

DK and VC conceived the project. DK prepared samples, collected and processed cryo-EM data, refined structures, and performed functional assays. PS prepared the sample for GPR119-G_q_ complex. AR performed the receptor expression test and functional assays. JS assisted with functional assays. MR and VK performed and analyzed MD simulations. VC supervised the project. DK prepared the initial manuscript draft with scientific input from VC, and all authors reviewed and edited the manuscript.

## Data availability

Structure coordinates and cryo-EM density maps have been deposited in the Protein Data Bank (PDB) and in the Electron Microscopy Data Bank (EMDB), respectively, under the following accession codes. OEA-bound GPR119-G_s_-Nb35 complex: PDB entry 36QE, EMDB-77772 (composite map), EMDB-77768 (consensus map), and EMDB-77769 (receptor-focused map); OEA-bound GPR119-G_q_ complex: PDB entry 36QF, EMDB-77773 (composite map), EMDB-77770 (consensus map), and EMDB-77771 (receptor-focused map).

## Materials and Methods

### Construct and insect cell expression

This study employed the engineered GPR119 and a dominant negative human Gα_s_ (DNGα_s_) which were designed in our previous study(*38*). Briefly, the wild-type (WT) human GPR119 (1-335) (UniProt ID Q8TDV5) contained a hemagglutinin (HA) signal peptide, FLAG-tag epitope (DYKDDDK), and 10× His tag, followed by an HRV 3C protease cleavage site and a thermally stabilized bRIL(*17*) at the N-terminus while the LgBiT(*18*) and eGFP were fused to the C-terminus with an HRV 3C protease cleavage site between them. For GPR119-G_s_ complex, the DNGα_s_ construct in a pFastBac1 vector was generated by site-directed mutagenesis to incorporate eight mutations including S54N, G225A, E268A, N271K, K274D, R280K, T284D, and I285T to stabilize the interaction with the βγ subunits(*39*). For GPR119-miniG_q_ complexes, the miniGα_q_ was designed as previously reported(*20*), based on a miniGa_s_ skeleton(*40*) with an addition of 18 aminoacids from the Ga_i1_ N-terminus and including two dominant-negative mutations (G203A and A326S)(*41*) to improve the stability of the GPCR-G_q_ protein complex. The human G_β1_ was fused with a 10× His tag and HRV 3C site at the N-terminus and with a 15aa-linker and HiBiT(*18*) at the C-terminus and cloned into a pFastBac dual vector together with the WT human G_γ2_.

The chimeric GPR119, DNGα_s_ or miniGα_q_, and G_β1γ2_ were co-expressed in *Sf9* insect cells using the Bac-to-Bac Baculovirus Expression System (Invitrogen). Cells were co-infected at a density of 2.0 × 10^6^ cells/mL with three separate viruses at a ratio of 1:1:1 for GPR119, DNGαs or miniGα_q_, and G_β1γ2_, respectively. The infected cells were incubated at 27 °C, harvested 48 h post-infection, washed with Phosphate-Buffered Saline (PBS), and stored at −80 °C until use.

### Expression and purification of Nb35

Nanobody 35 (Nb35)(*21*) expressing cells were cultured as described in our previous study(*42*). Cells were thawed and resuspended in TES Buffer (50 mM Tris pH 8.0, 20% (w/v) sucrose, 0.5 mM EDTA). After 1.5 h of stirring, two equal volumes of the 4× diluted TES were added to initiate hypotonic lysis. After an additional 1 h of stirring, the soluble supernatant was subjected to a HisPur™ Ni-NTA Resin (Thermo Scientific). The eluted Nb35 was concentrated and injected into a Superde× 10/300 GL column (Cytiva) pre-equilibrated with 20 mM HEPES pH 7.5, 100 mM NaCl, and a monodisperse fraction of Nb35 was collected and concentrated to 10 mg/mL using a 10-kDa cutoff Amicon Ultra Centrifugal Filter (Millipore). The final buffer was exchanged to 20 mM HEPES pH 7.5, 150 mM NaCl, 20% glycerol, and flash-frozen in aliquots, stored at −80 °C for future use.

### Purification of the OEA-bound GPR119-G_s_-Nb35 and GPR119-miniG_q_ complexes

The target protein expressing *Sf9* cells were lysed in a buffer containing 20 mM HEPES, pH 7.5, 100 mM NaCl, 5 mM MgCl_2_, 5 mM CaCl_2_, 0.1 mM TCEP, and 10% (v/v) glycerol, supplemented with a homemade protease inhibitor cocktail (500 µM AEBSF, 1 µM E-64, 1 µM leupeptin, and 150 nM aprotinin). Both GPR119-G_s_ and G_q_ complex were assembled by adding 100 μM OEA (Cayman company), 25 mU/mL apyrase, and 10 μg/mL Nb35 for only G_s_ complex, followed by 1 h incubation at room temperature. The membrane fraction was collected by centrifugation at 30,000×g for 30 min and then was solubilized in 20 mM HEPES, pH 7.5, 100 mM NaCl, 5 mM MgCl_2_, 5 mM CaCl_2_, 0.1 mM TCEP, 0.5% (w/v) lauryl maltose neopentyl glycol (LMNG, Anatrace), 0.05% (w/v) cholesterol hemisuccinate (CHS, Anatrace), 25 mU/mL apyrase, and 100 μM OEA at 4 °C for 2 h. The supernatant was isolated by centrifugation at 30,000×g for 30 min and was further incubated overnight at 4 °C with an anti-GFP nanobody resin, which was made of AminoLink Resin (Thermo Scientific) coupled with anti-GFP nanobody(*43*). The resin was washed with 20 column volumes (cv) of 20 mM HEPES, pH 7.5, 100 mM NaCl, 10% (v/v) glycerol, 0.1 mM TCEP, 0.1% (w/v) LMNG, 0.01% (w/v) CHS, 100 μM OEA and additional 20 cv of the same buffer containing 0.05% (w/v) LMNG, 0.005% (w/v) CHS. Then the resin was incubated with 5 cv of Buffer containing 0.02% (w/v) LMNG, 0.002% (w/v) CHS with HRV 3C protease (GenScript) at 4 °C for 3 h. The eluted sample was injected into a Superose 6 Increase 10/300 column (Cytiva) for the G_s_ complex or Superdex 200 GL 10/300 column (Cytiva) for the G_q_ complex, each pre-equilibrated with an SEC buffer containing 20 mM HEPES pH 7.5, 100 mM NaCl, 0.1 mM TCEP, 0.00075% (w/v) LMNG, 0.000075% (w/v) CHS, and 100 μM OEA. A monodisperse peak fraction corresponding to the complex was pooled and further concentrated to 2∼3 mg/mL for cryo-EM, using a 100-kDa cutoff Amicon Ultra Centrifugal Filter (Millipore).

### Cryo-EM grid preparation

Cryo-EM grids were prepared by applying 3 μl samples onto glow-discharged Quantifoil holey carbon grids (Au, R1.2/1.3, 200 mesh) at 20 mA for 40 s and blotting for 3 s, followed by plunge-freezing into liquid ethane using a Vitrobot Mark IV (Thermo Scientific) operating at 100% humidity and 4 °C.

### Cryo-EM data acquisition and image processing

Cryo-EM data were collected on a Titan Krios (Thermo Fisher Scientific) at the Core Center of Excellence in Nano Imaging (CNI) of the University of Southern California, equipped with a Gatan BioQuantum K3 Imaging Filter and a Gatan K3 direct-electron detector and operated at an acceleration voltage of 300 kV. Images were recorded at a nominal magnification of 105,000 x resulting in the pixel size of 0.85 Å. A total of 8,040 and 11,199 movies were recorded for OEA-bound GPR119-G_s_ and -G_q_ complexes, respectively, with a defocus range of −0.8 to −2.4 µm and a total dose of 60 e^−^/A^2^.

The overall cryo-EM data processing workflow is shown in Supplementary Fig. S1 and S2. Both datasets were processed with cryoSPARC (v4.7.1)(*44*) unless stated otherwise. Cryo-EM movie stacks were aligned using Patch motion correction followed by Patch CTF estimation. Micrographs were curated using a CTF resolution cut off < 4.0 Å. A total of 6,300 and 11,026 micrographs for G_s_ and G_q_ complex structures, respectively, were selected, in which 500 or 1,000 micrographs from each dataset were used for the blob-picker and iterative 2D classification. The selected particles were then used as a reference for template picking and the resulting particles were used for Topaz train, followed by Topaz extraction with the entire micrographs. After particle extraction with box size of 324 pixels (Fourier cropped to 64 pix) and 2D classification, 710,476 and 2,913,752 particles were selected for the OEA-bound GPR119-G_s_ and -G_q_ complexes, respectively, followed by *ab-initio* 3D reconstruction and heterogeneous refinement. The 358,096 and 599,088 particles selected from heterogeneous refinement for each dataset were re-extracted with box size of 324 pixels. Subsequently, the extracted particles underwent 2D classification to get rid of the residual junk particles. Next, a non-uniform refinement and following local refinements with masks focused on the receptor further improved the map quality. The final composite maps were generated by Combine Focused Maps in Phenix (v.1.21)(*45*) using consensus map and receptor-focused map, resulting in gold-standard Fourier shell correlation (GSFSC) resolutions of 2.72 Å and 3.05 Å for the OEA-GPR119-G_s_ and -G_q_ complex, respectively. All maps were sharpened by Sharpening Tool in cryoSPARC (v.4.7.1)(*46*).

### Model building and refinement

The initial GPR119-G_s_-Nb35 model (PDB ID 8VHF)(*38*) was fit into the obtained cryo-EM map by ChimeraX(*47*). The model was then subject to iterative cycles of Real-Space Refine in Phenix(*48*) and manual inspection and building in COOT(*49*). Structural figures were created by ChimeraX(*47*). Data collection and refinement statistics are shown in Supplementary Table S1.

### Cell culture, transfection, and constructs for functional assays

Human Embryonic Kidney 293T (HEK293T) cells (ATCC) were maintained, passaged, and transfected in culture flasks using Complete Dulbecco’s Modified Eagle Medium (DMEM) Media (Thermo Fisher Scientific) supplemented with 10% Fetal Bovine Serum (FBS) (Cytiva), 2 mM L-Glutamine (Thermo Fisher Scientific), 100 U/mL-100 µg/mL Penicillin-Streptomycin (Thermo Fisher Scientific) in a humidified atmosphere at 37°C and 10% CO_2_. Cells were dissociated using 0.05% trypsin-EDTA (Thermo Fisher Scientific) and passed every 3-4 days at less than 90% confluence.

In this study, otherwise noted, the cells were seeded in a 6-well plate (Corning) at a density of 350,000 cells per well in 2 mL of complete DMEM and incubated overnight. The following day, cells were transfected with respective plasmids and empty vector pcDNA3.1 in total 1000 ng using polyethyleneimine (PEI) (PEI-STAR, Tocris) at a ratio of 1:3 DNA(µg):PEI(µg). After 24 h post-transfection, the cells were collected and the assays were performed.

For functional assay, the WT human GPR119 (1-335) (UniProt ID Q8TDV5) gene sequence contained a hemagglutinin (HA) signal peptide and FLAG-tag epitope at the N-terminus was cloned into a pCDNA3.1 Vector. For surface expression test, 4xGSA linker was inserted between the FLAG-tag and GPR119 to facilitate the access of Anti-Flag antibody on the membrane surface. Mutations were introduced to this WT GPR119 construct by site-directed mutagenesis and the sequences for all constructs were verified using commercial service (Genewiz).

### BRET-based G_s_ protein coupling assay (ONE-GO)

The G_s_ signaling was measured using ONE-GO assay(*50, 51*). Briefly, this assay directly measures G protein activation by monitoring Bioluminescence Resonance Energy Transfer (BRET) between a selective biosensor and the GTP-bound Gα subunit. The biosensor comprises a single plasmid encoding a specific YFP-fused Gα subunit and a membrane-anchored nanoluciferase (NLuc) fused to a peptide or protein domain exhibiting high selectivity binding towards the corresponding Gα-GTP complex. Upon receptor activation, the cognate GDP-bound G proteins undergo conformational transformations resulting in the exchange of GDP to GTP and subsequent dissociation of Gα-GTP and free Gβγ from the receptor. An increase in the BRET ratio between the Nluc-tagged detector and YFP-tagged Gα subunit provides a direct measure of receptor signaling through its cognate G proteins.

HEK293T cells were transfected with 10 ng of WT GPR119 or mutants in pcDNA3.1, 50 ng of ONE-GO Gα_s_ sensor, and 940 ng of empty vector pcDNA3.1 using PEI as described above. In the following day, the cells were detached and resuspended into BRET buffer (140 mM NaCl, 5 mM KCl, 1 mM MgCl_2_, 1 mM CaCl_2_, 0.37 mM NaH_2_PO_4_, 20 mM HEPES, 0.1% Glucose). To carry out the assay, 300× OEA (30 mM to 30 μM) (Cayman Chemical) and 200× coelenterazine 400a (CTZ400a) (2 mM) (Nanolight Technologies) stocks were prepared in DMSO and Nanofuel solution (Nanolight Technologies), respectively. Both stock solutions were diluted 100 times in the assay buffer to achieve 3× of the final concentration. 15 µL of the graded concentration of 3× OEA was transferred into corresponding wells of a white bottom 384-well plate (Revvity) and 15 µL of 3× CTZ400a was added into each well, followed by pulse spin-down at 200× *g*. Finally, 15 µL of cells were added into each well at a density of ∼15,000 cells/well, followed by pulse spin-down at 200× *g*. The BRET2 signal was measured with a PHERAstar FSX plate reader (BMG Labtech) at 28 °C using 0.32 s integration time and BRET2 filters (460/30 nm and 535/30 nm) starting 5 min after the addition of cells. The netBRET signal was calculated as the ratio between the emission intensity at 535 nm divided by the emission intensity at 460 nm. ΔBRET was calculated as the difference between the netBRET signals collected in the presence and absence of the ligand. The ΔBRET values for each mutant were normalized to the maximal ΔBRET signal of WT (100%). BRET data were collected from three biologically independent samples in triplicates and analyzed using non-linear regression with a log(ligand) vs response (3-parameter) function in GraphPad Prism v. 11.0.2.

### Inositol monophosphate IP-ONE accumulation assay

Gα_q_-mediated inositol monophosphate (IP_1_) accumulation was measured using the homogeneous time-resolved fluorescence (HTRF) IP-ONE G_q_ assay Kit (Revvity). Briefly, the assay is based on a competitive time-resolved Förster resonance energy transfer (TR-FRET) principle. Upon receptor activation, endogenous IP_1_ competes with acceptor-labeled IP_1_ (IP_1_-d2) for binding to donor-labeled anti-IP_1_ antibodies (Tb cryptate), displacing IP_1_-d2 and thereby reducing the basal FRET signal. The decrease in TR-FRET intensity is inversely proportional to the amount of IP_1_ produced, providing a sensitive and quantitative measure of receptor activation. HEK293T cells were seeded into 6-well plates at a density of 350,000 cells per well and after 24 h incubation, the media was replaced with 2 mL of DMEM containing 1% FBS and 400 ng of WT GPR119 or mutant DNAs were transfected using PEI as described above. After 24 h post-transfection, cells were detached and resuspended into 1x Stimulation buffer, and 7 µL of cells were added into low volume 384-well white plates (Corning) with ∼15,000 cells per well. The cells were stimulated with 7 µL of the graded concentration of OEA for 2 h at 37 °C and subjected to incubation with each 3 µL of IP_1_-d2 and IP1 Tb Cryptate Antibody for 1.5 h with shaking at 500 rpm. The FRET signal was measured using a PHERAstar FSX plate reader (BMG Labtech) at 22 °C. The netFRET signal was calculated as the ratio between the emission intensity at 665 nm (acceptor) divided by the emission intensity at 620 nm (donor). ΔFRET was calculated as the difference between the netFRET signals collected in the presence and absence of the ligand. The ΔFRET values for each mutant were normalized to the minimal value of ΔFRET of WT (100%). FRET data were collected from three biologically independent samples in duplicates or triplicates and analyzed using non-linear regression with a log(ligand) vs response (3-parameter) function in GraphPad Prism v. 11.0.2.

### Surface expression measurement by flow cytometry

Surface expression levels of the WT GPR119 and its mutants were measured using flow cytometry by detecting the N-terminal FLAG (DYKDDDK) tag. A 1 mg/mL ANTI-FLAG M2-FITC antibody stock (Sigma Aldrich) was diluted by 100x using 1× TBS (Corning) supplemented with 4% Bovine Serum Albumin (BSA) (Sigma Aldrich). A commercial 7-AAD stock (50 µg/mL) (Thermo Fisher Scientific) was then supplemented to the prepared Anti-FLAG M2-FITC solution to achieve final concentrations of 8.3 µg/mL for both ANTI-FLAG M2-FITC and 7-AAD.

HEK293T cells were seeded in a 12-well plate (Corning) at a density of 140,000 cells per chamber in 800 µL of Complete DMEM Media and incubated overnight. After 24 h, the cells were transfected with 400 ng of a given receptor construct (including empty vector as a negative control) using PEI as described above. Forty-eight hours after transfection, the media was aspirated, and the cells were detached via trituration using the BRET buffer, centrifuged, and resuspeneded in 20-30 µL of BRET buffer. A 10 µL aliquot of cells for each construct was then mixed with 10 µL of M2-FITC + 7-AAD working stock solutions prepared above. The plate was incubated for 20 min at 4 °C, followed by dilution with 180 µL of 1x TBS buffer (Corning). Expression levels were then measured using a Millipore Guava EasyCyte™ 6HT flow cytometer controlled by the Millipore guavaSoft - ExpressPlus (v.3.6) software, with the instrument being set to a gated count of 10,000 live cells using Red fluorescence (640 nm laser and 680/30 nm fluorescence filter) vs forward scattering plot. Surface expression was measured using Green fluorescence (488 nm laser and 525/30 nm fluorescence filter). Instrument gain and fluorescence threshold settings were established using WT GPR119- and empty vector-transfected cells to achieve optimal separation between positive and negative cell populations (Supplementary Fig. S17). To determine the relative surface expression of each construct, the median Green fluorescence signal of each construct in the Green-positive/Red-negative quadrant was normalized by the median green fluorescence signal of the WT receptor. Surface expression levels were obtained from three biologically independent samples. Expression of GPR119 mutants was compared with that of the WT receptor using a one-way mixed-effects model with biological experiment as a blocking factor, followed by Dunnett’s multiple-comparisons test in GraphPad Prism v. 11.0.2.

### Molecular Dynamics Simulations

GROMACS v2026.0(*52*) was used for all simulations. The CHARMM36 all-atom(*53*) force field was employed for parameterization and topology generation. The CHARMM-GUI web server was used to generate system parameters and topologies(*54–56*).

Two initial conformations for OEA-GPR119-G_s_ and OEA-GPR119-G_q_ complexes were obtained from cryo-EM studies. Two additional models were constructed based on these structures as templates, in which dominant-negative Gα_s_ and miniGα_q_ were replaced with wild-type Gα_s_ and Gα_q_, respectively. All models were independently embedded in a lipid bilayer composed of 100% 1-palmitoyl-2-oleoyl-sn-glycero-3-phosphocholine (POPC), following previous GPCR simulation studies(*57, 58*). To maintain consistency across systems, the α-helical domain (AHD) of Gα, which was unresolved in the cryo-EM structures, was not modeled in any system.

Missing residues were modeled as follows. In all systems, the C-terminal region of Gγ (residues 63–71) was added. In the GPR119–DNGα_s_ system, receptor residues 212–218, as well as DNGα_s_ N-terminal residues 2–10 and residues 255–261, were modeled. The same regions were added in the GPR119–WT Gα_s_ system. In the GPR119–miniGα_q_ system, receptor residues 190–218 and miniGα_q_ residues 2–5 and 64–66 were modeled. In the GPR119–WT Gα_q_ system, missing receptor residues were modeled, and miniGα_q_ was replaced with WT Gα_q_; residues 1–7 of Gα_q_ were not modeled, and the AHD was omitted as described above.

In all systems, Cys68 of Gγ was geranylgeranylated. Cys3 of DNGα_s_, WTGα_s_, and miniGα_q_, as well as Cys9 of WTGα_q_, were palmitoylated. The C-terminus of Gγ was capped with a methylamide group. Membrane positioning was performed using the Positioning of Proteins in Membranes (PPM) server via the CHARMM-GUI interface. All systems were solvated with TIP3P water molecules. Sodium and chloride ions were placed to achieve a final concentration of 150 mM NaCl. Based on system size, the membrane plane dimensions (XY) were set to 120 Å in a hexagonal box, and the Z dimension was set to 160 Å.

Energy minimization was performed using the steepest descent algorithm for 50,000 steps with a convergence criterion of 100 kJ·mol⁻¹·nm⁻¹. This was followed by six equilibration steps totaling 10 ns, conducted under NVT and subsequently NPT ensembles with a time step of 1–2 fs, using a V-rescale thermostat at 303.15 K and a C-rescale barostat at 1 atm. In the final equilibration step, the barostat was switched to Parrinello–Rahman.

For each system, ten independent production runs of 1 μs were performed under the NPT ensemble using the Parrinello–Rahman barostat (1 atm) and V-rescale thermostat (303.15 K), with randomized initial velocities. Simulations were executed on GPU clusters at the Center for Advanced Computing at the University of Southern California.

For analyzing the MD trajectories, GROMACS analysis toolkit, the MDTraj software package(*59*) and MDCiao(*60*) were used.

### Statistical Analysis

All statistical analyses were performed using Prism v. 11.0.2. (GraphPad Software Inc). Multiple comparisons between WT and mutants were carried out using one-way ANOVA followed by Dunnett’s multiple comparisons test. Significant differences are displayed as *p<0.05, **p<0.01, ***p<0.001, and ****p<0.0001. All the data were from at least three biologically independent experiments and values are presented as means ± SEM.

## Supplementary Information

**Supplementary Figure S1.**
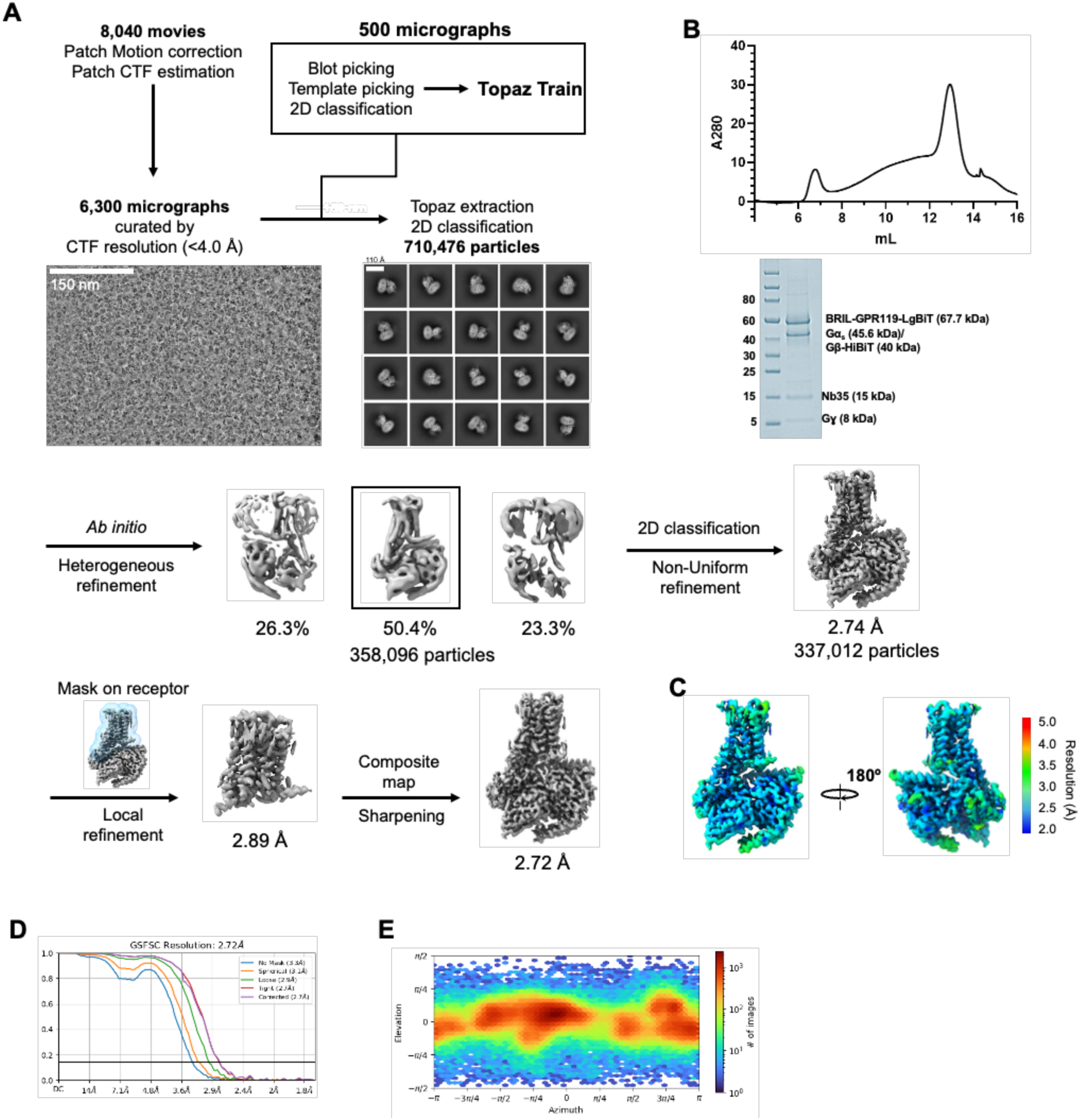
Cryo-EM data processing for OEA-bound GPR119-G_s_-Nb35 complex. (**A**) Cryo-EM workflow for OEA-bound GPR119-G_s_-Nb35 complex. (**B**) Representative size-exclusion chromatography (SEC) elution profile and Coomassie-stained SDS-PAGE gel of purified OEA-bound GPR119-G_s_-Nb35 complex. (**C**) Local resolution map in two different orientations. (**D**) Gold-Standard Fourier Shell Correlation (GSFSC) curve. (**E**) Angular distribution of particles used for the final map.

**Supplementary Figure S2.**
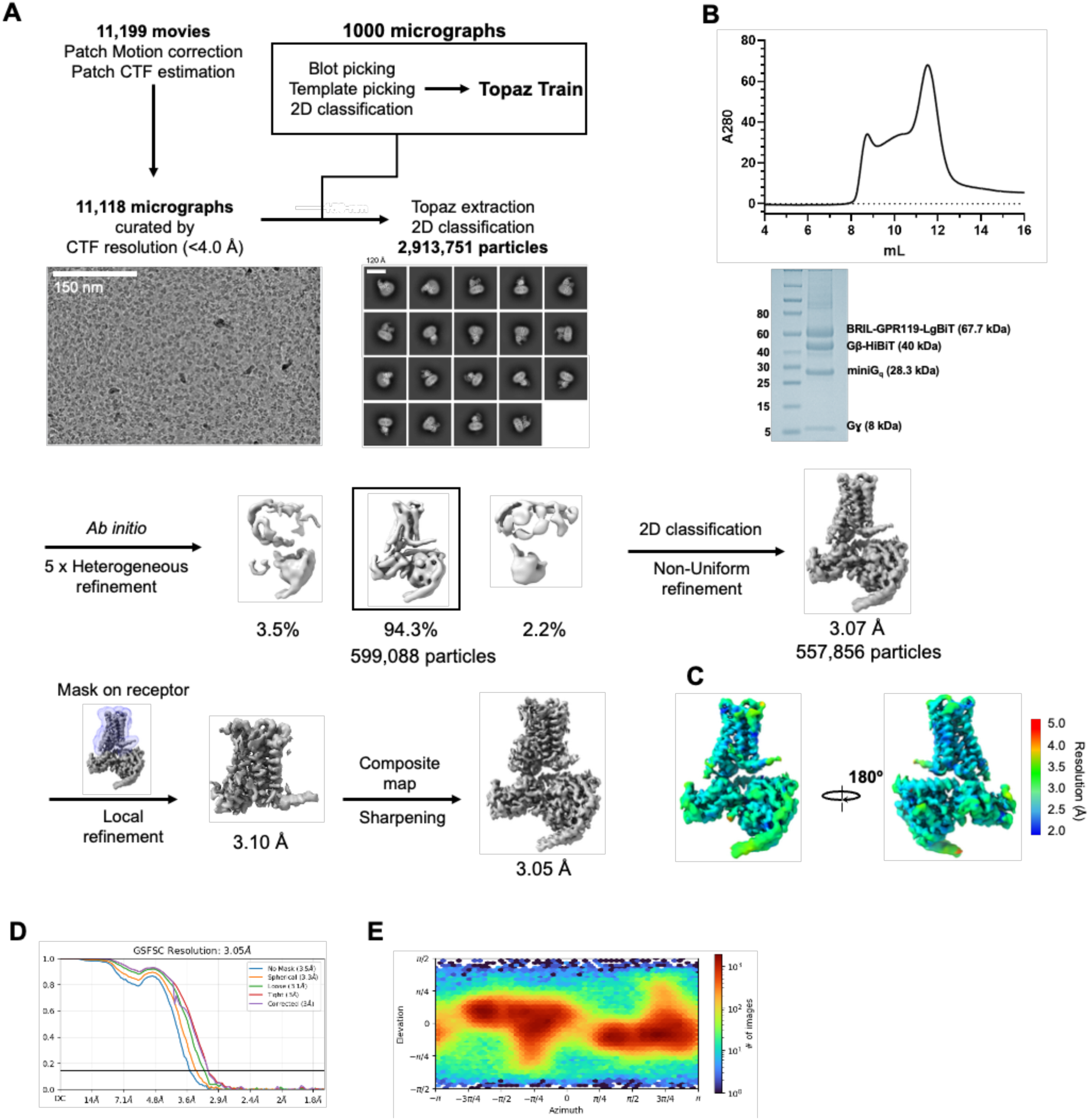
Cryo-EM data processing for OEA-bound GPR119-G_q_ complex. (**A**) Cryo-EM workflow for OEA-bound GPR119-G_q_ complex. (**B**) Representative size-exclusion chromatography (SEC) elution profile and Coomassie-stained SDS-PAGE gel of purified OEA-bound GPR119-G_q_ complex. (**C**) Local resolution map in two different orientations. (**D**) Gold-Standard Fourier Shell Correlation (GSFSC) curve. (**E**) Angular distribution of particles used for the final map.

**Supplementary Figure S3.**
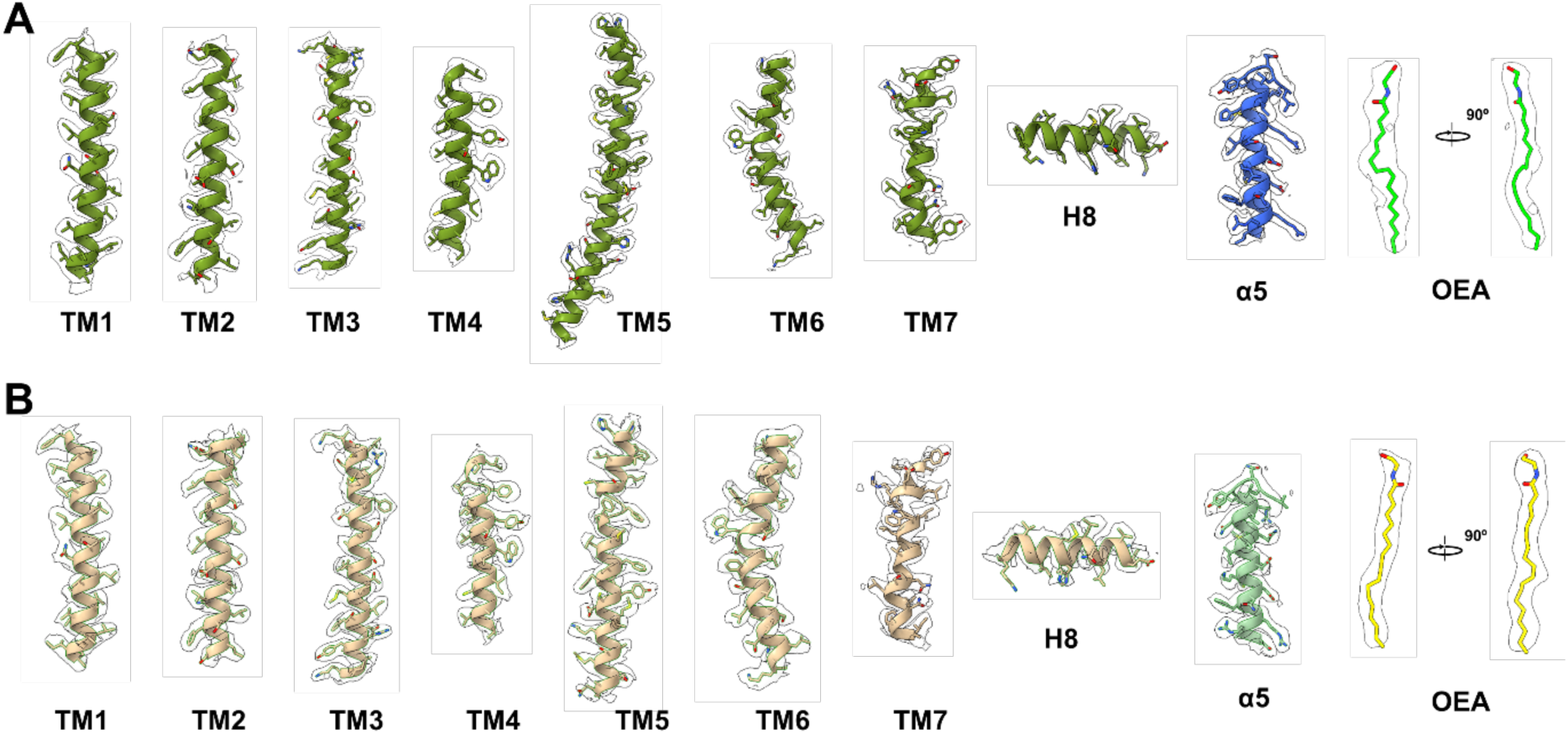
Cryo-EM density maps of representative regions of GPR119-G protein complexes. (**A**) EM density maps of each TM helix of GPR119, α5 helix of Gα_s_, and OEA in OEA-bound GPR119-G_s_ complex at a contour level 7. (**B**) EM density maps of each TM helix of GPR119, α5 helix of Gα_q_, and OEA in OEA-bound GPR119-G_q_ complex at a contour level 9.

**Supplementary Figure S4.**
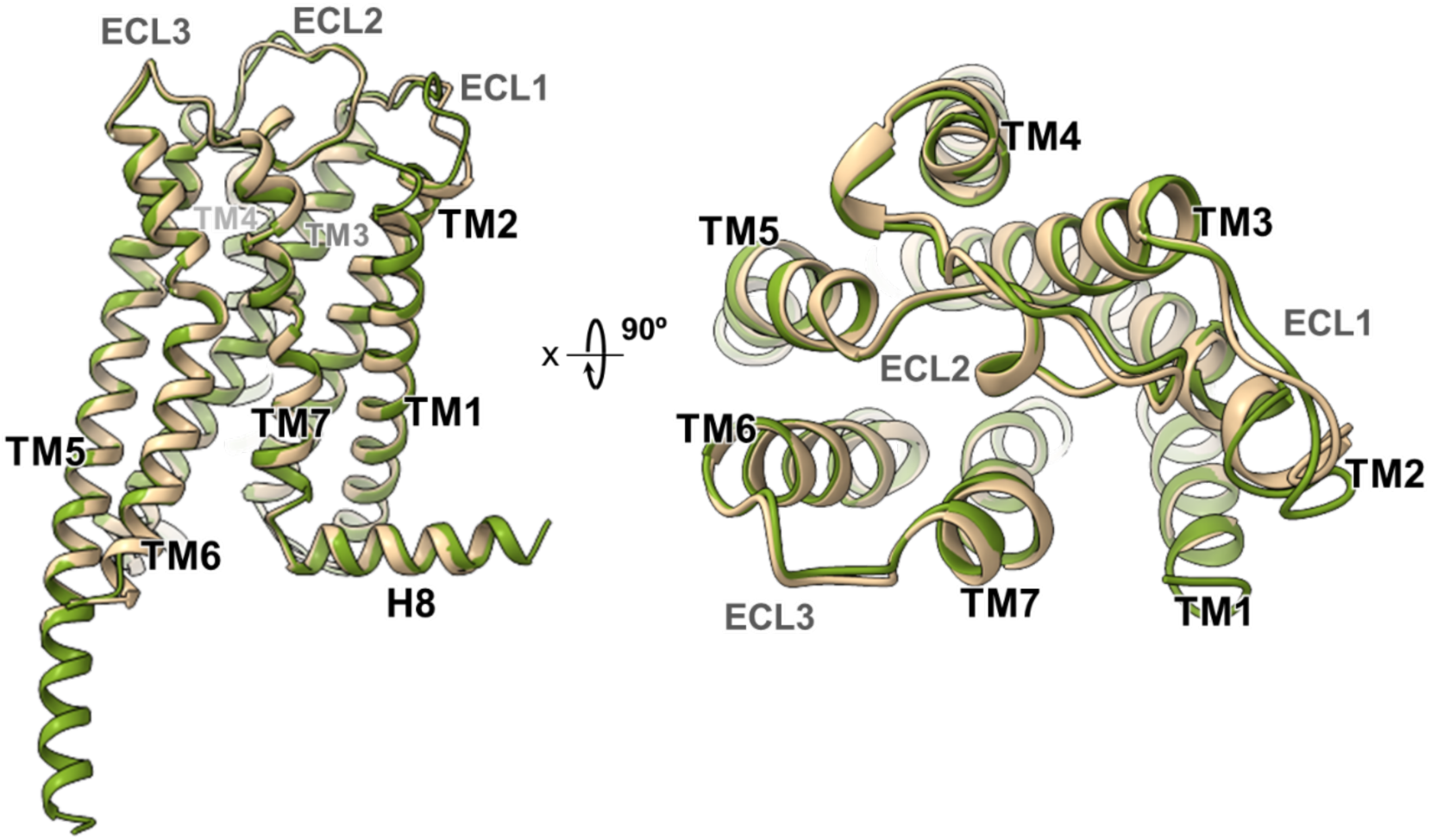
Comparison of GPR119 structures in G_s_- and G_q_-coupled complexes. GPR119 models from G_s_-coupled (green) and G_q_-coupled (pale) complexes are superimposed and shown in two different views.

**Supplementary Figure S5.**
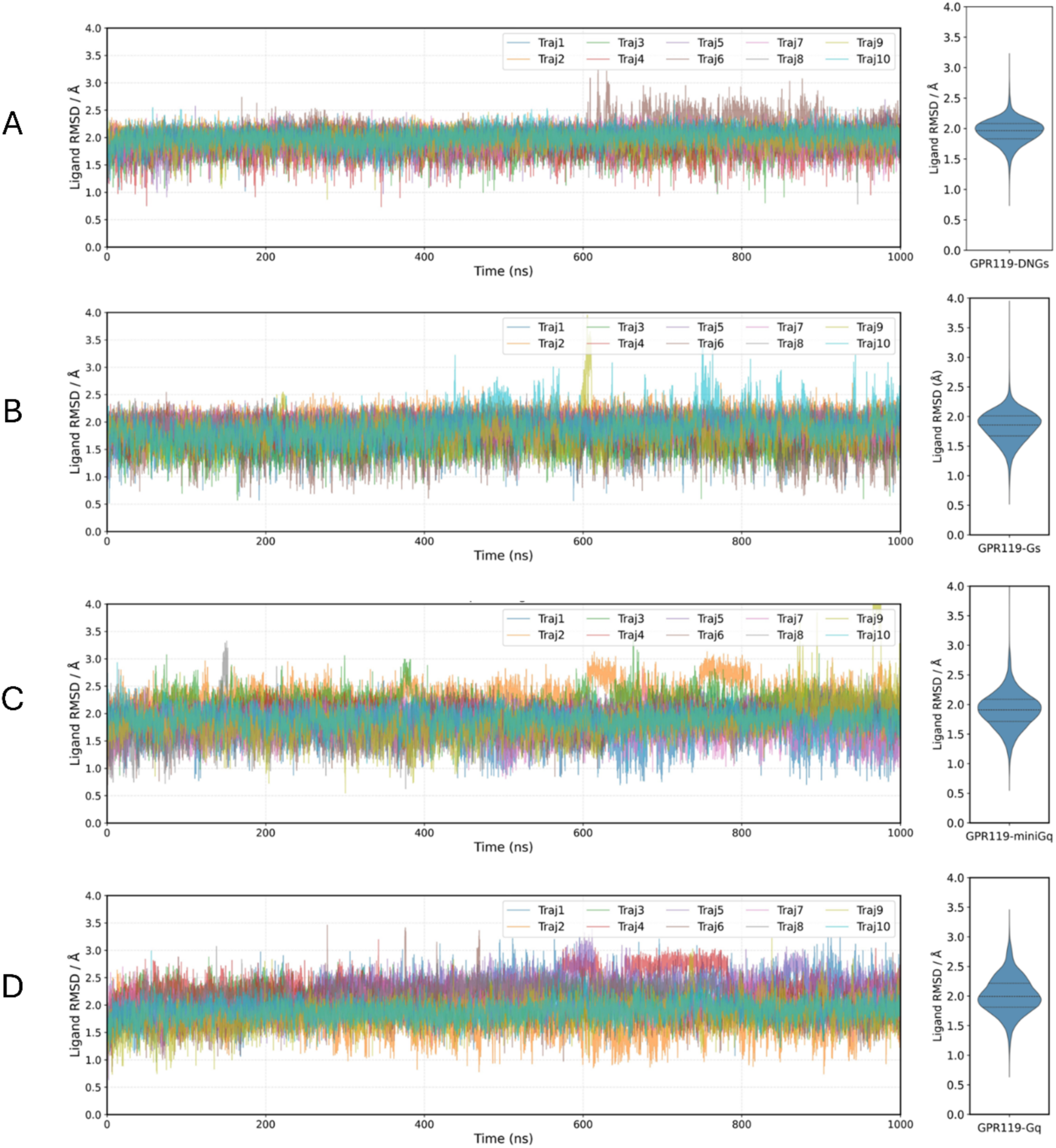
Traces of ligand RMSD for all MD trajectories of each complex. **(A**) GPR119-DNG_s_, (**B**) GPR119-G_s_, (**C**) GPR119-miniG_q_, (**D**) GPR119-G_q_. On the right side of each trace, violin plots show distributions of the ligand RMSD for each structure.

**Supplementary Figure S6.**
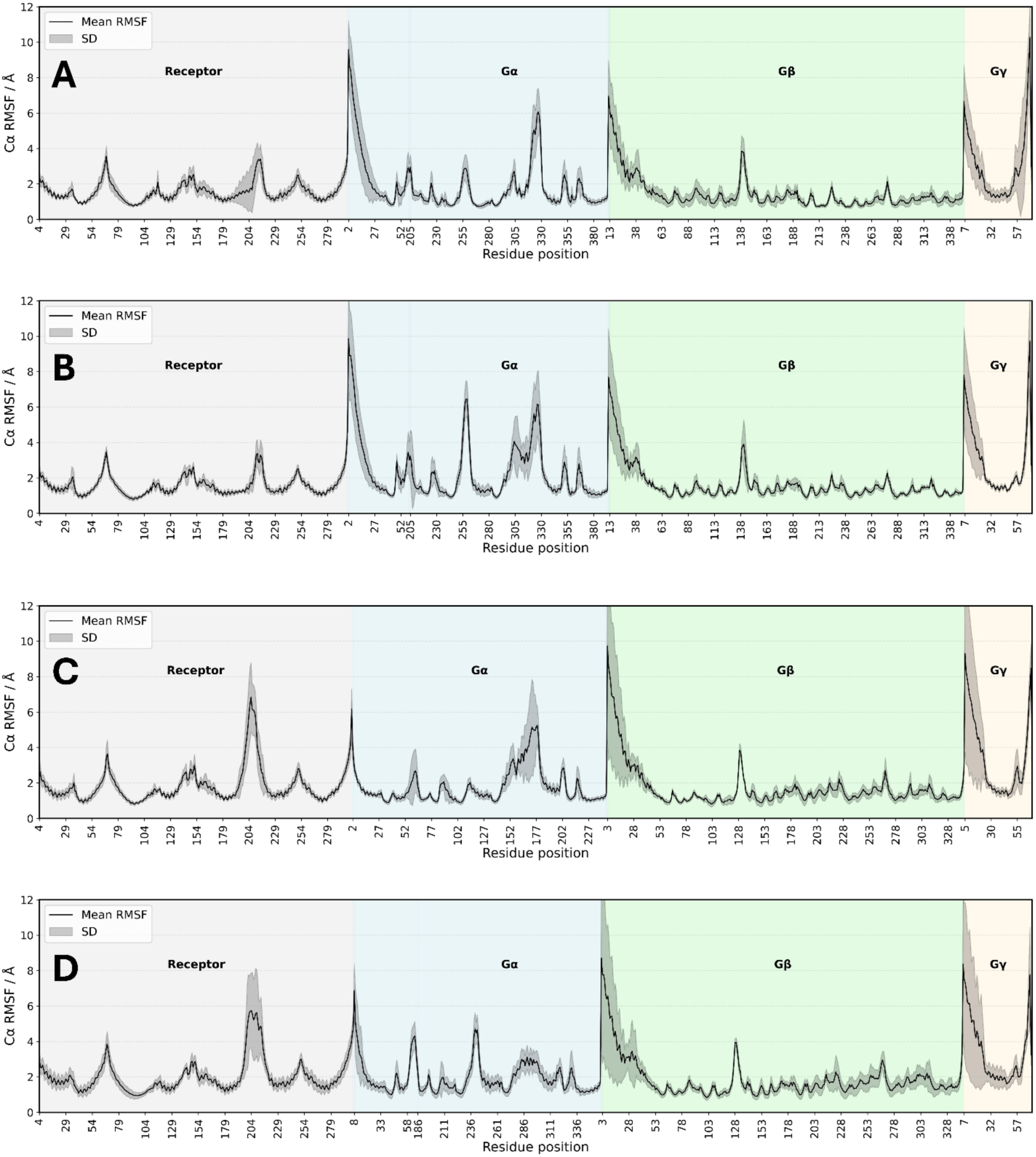
Comparison of Root Mean Square Fluctuation (RMSF) of all residues in MD simulations. **(A**) GPR119-DNG_s_, (**B**) GPR119-G_s_, (**C**) GPR119-miniG_q_, (**D**) GPR119-G_q_.

**Supplementary Figure S7.**
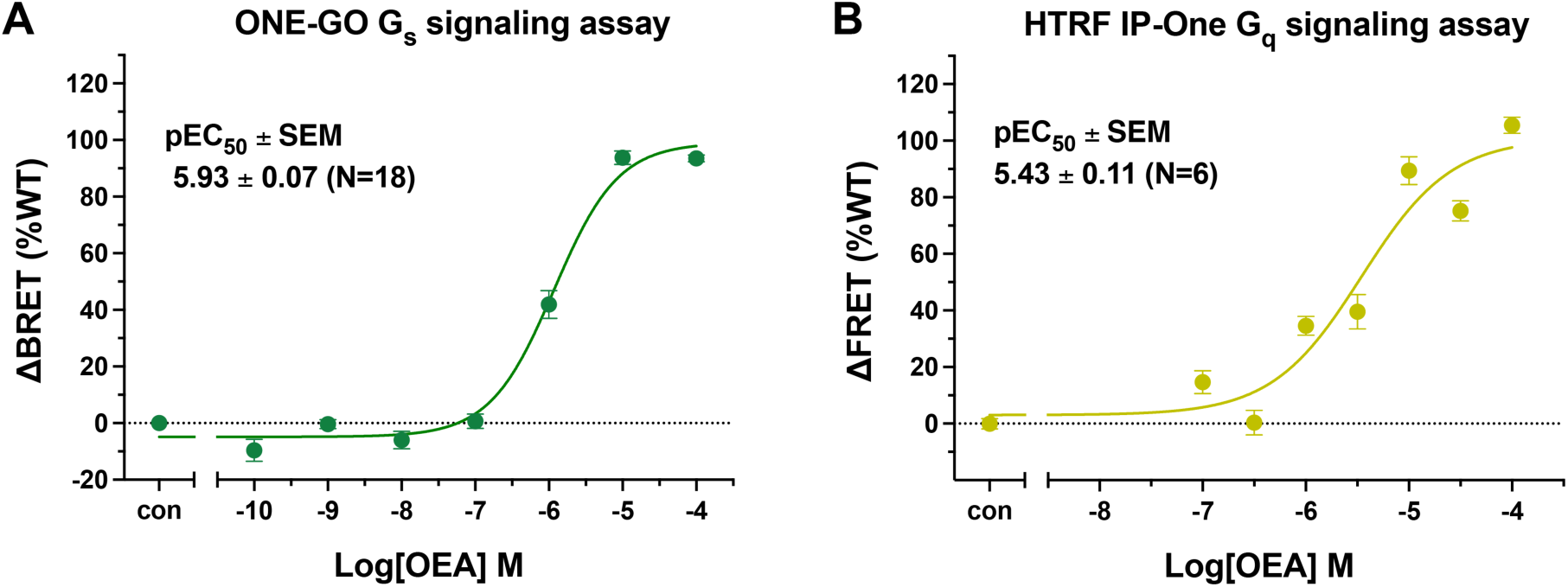
Pharmacological characteristics of OEA assessed in G_s_ and G_q_ signaling assay. (**A**) OEA-induced G_s_ signaling measured by ONE-GO G_s_ activation assay (**B**) OEA-induced G_q_ signaling measured by HTRF IP-One accumulation assay. Data points correspond to mean ± SEM from N=18 (A) or N=6 (B) independent experiments.

**Supplementary Figure S8.**
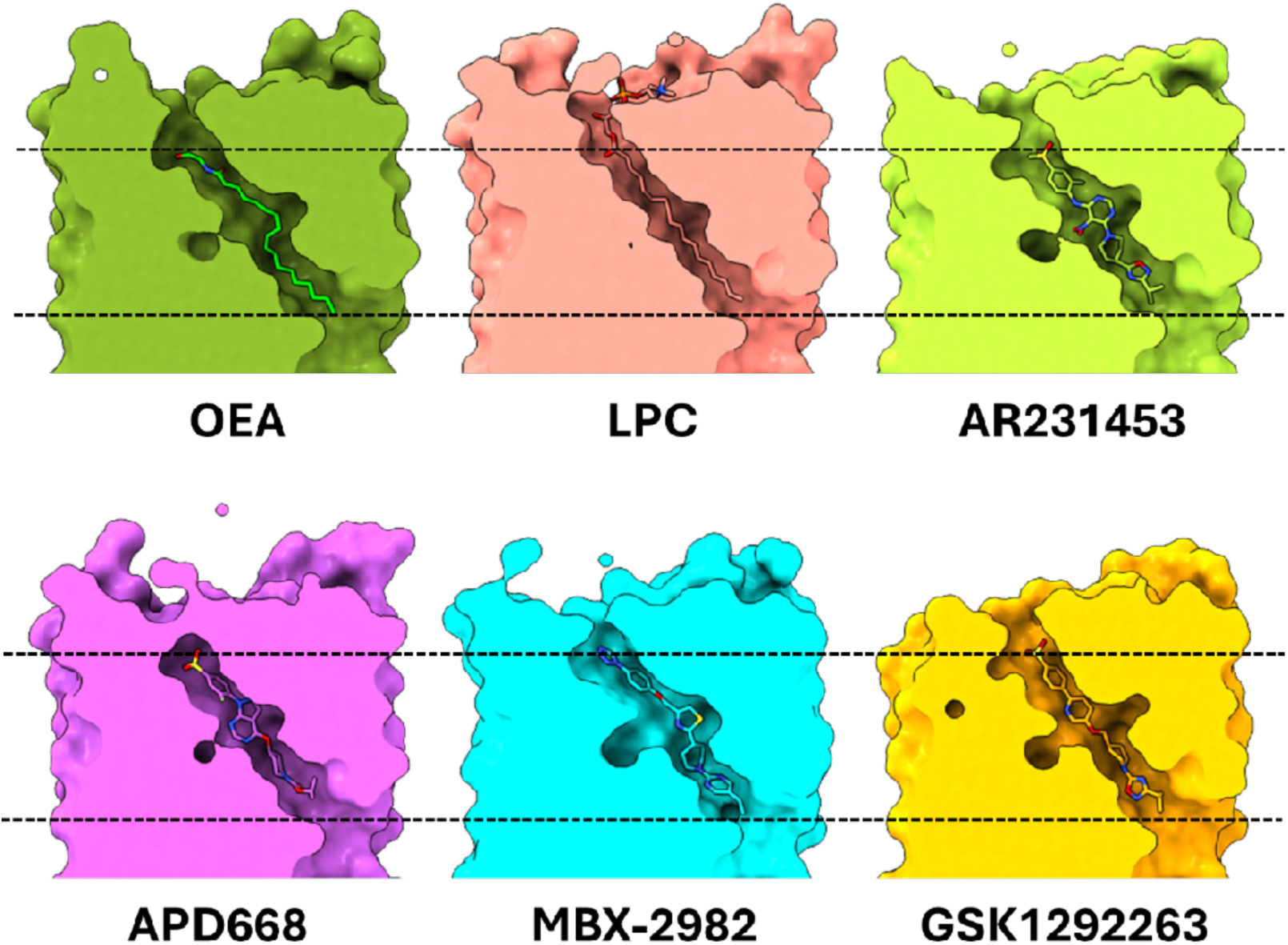
Comparison of the ligand-binding pocket in OEA-GPR119-Gs with previously reported GPR119-G_s_ structures. The ligand-binding pockets are shown as surface view, with bound ligands depicted as sticks within the transmembrane cavity. Dashed lines indicate the upper and lower boundaries of the OEA-binding pocket. The GPR119-G_s_ structures bound to LPC, AR231453, APD668, MBX-2982, and GSK1292263 were retrieved from the Protein Data Bank (PDB) under accession codes 7XZ5, 7WCN, 7XZ6, 7WCM, and 8ZRK, respectively.

**Supplementary Figure S9.**
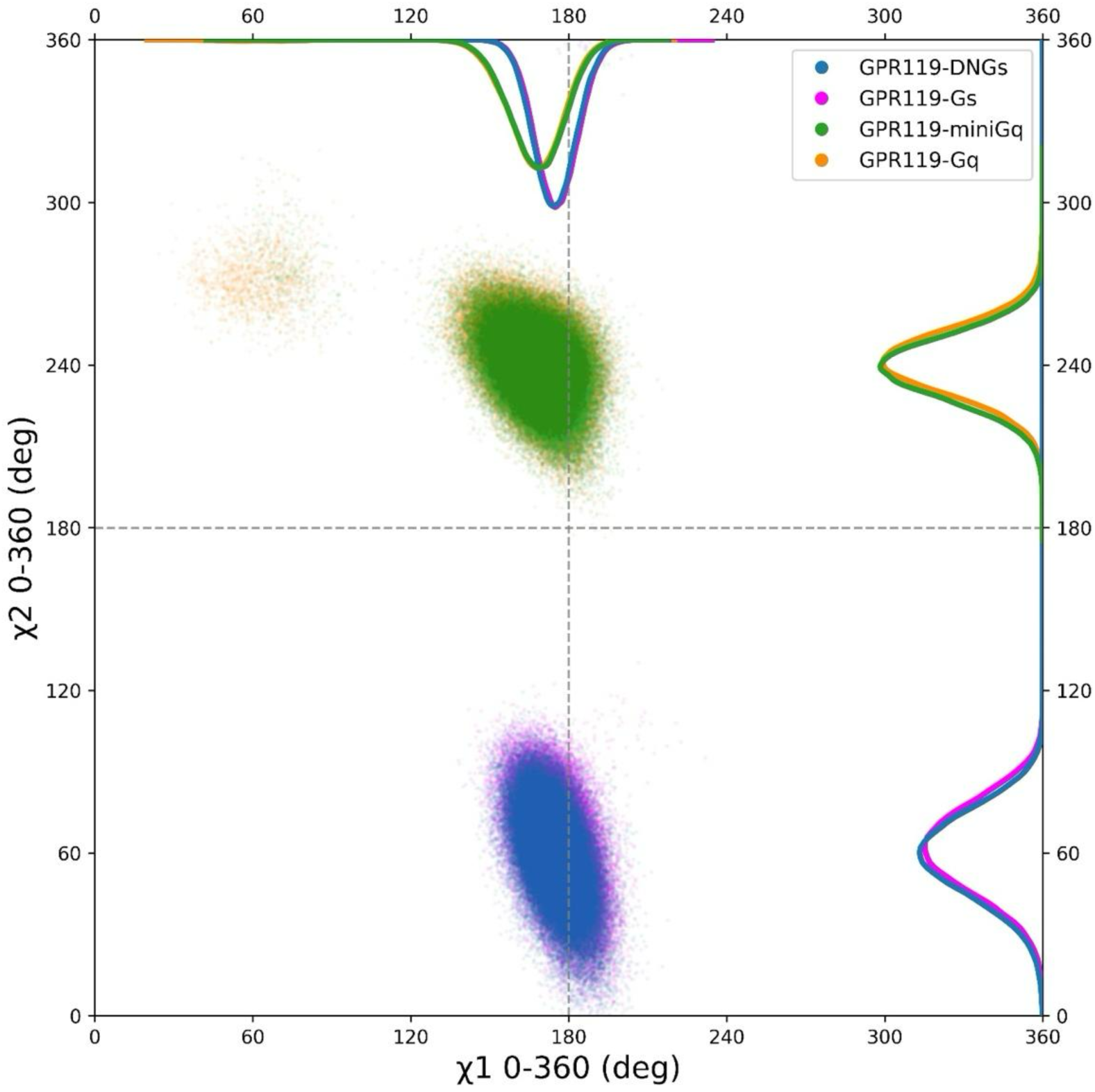
Comparison of side chain conformation of residue W265^7.39^ across all MD simulations. The X-axis represents the χ1 angle, and the Y-axis shows χ2.

**Supplementary Figure S10.**
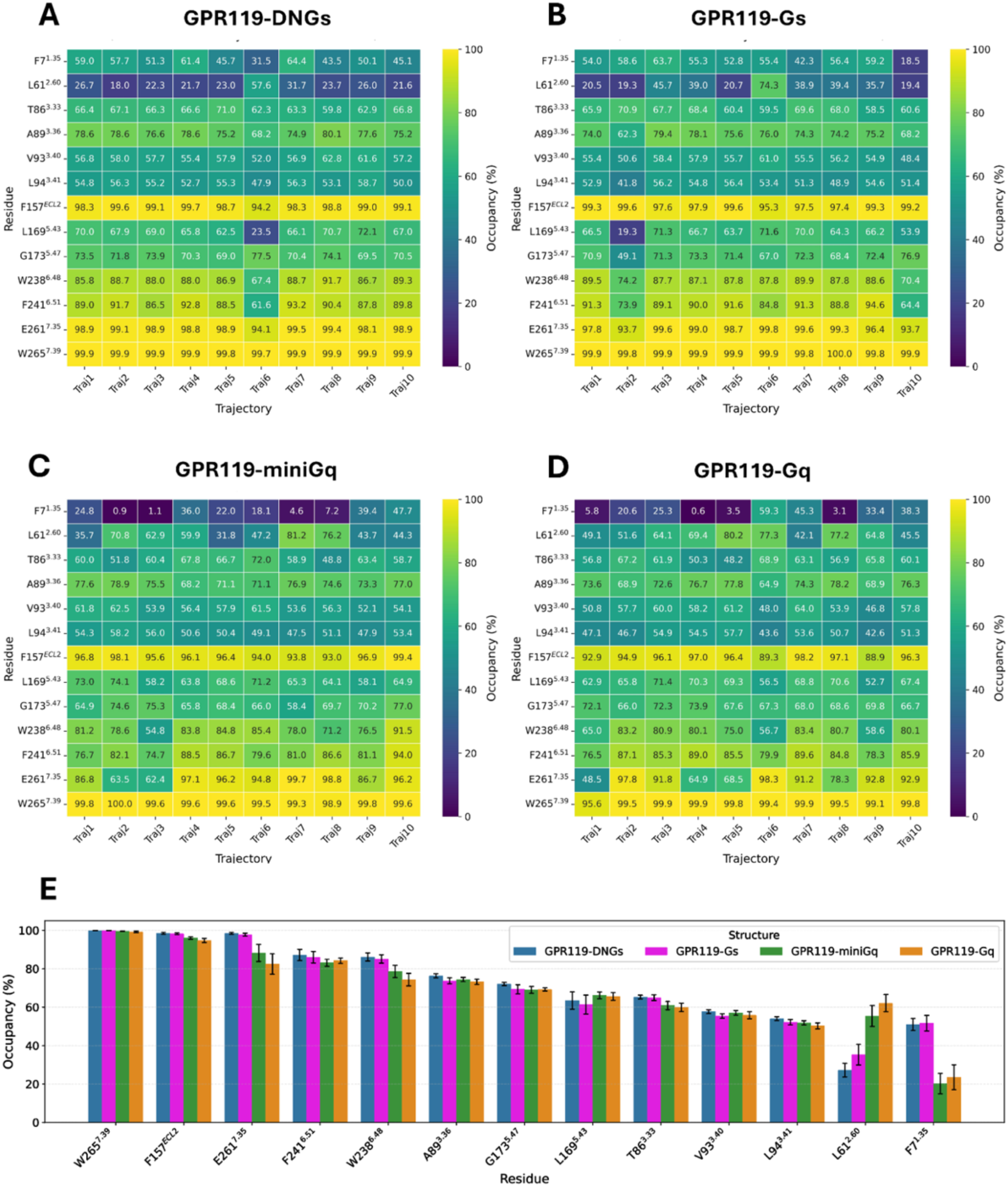
MD analysis of the receptor-ligand interactions. **(A-D)** Heatmaps showing the contact occupancy between OEA and the binding pocket residues (defined as the percentage of simulation frames where the closest heavy-atom distance between OEA and residue is ≤ 4 Å) across all trajectories. **(E)** Ligand pocket contact occupancy across structures. Colored bars show the average of all trajectories per structure with error bars representing ±SEM.

**Supplementary Figure S11.**
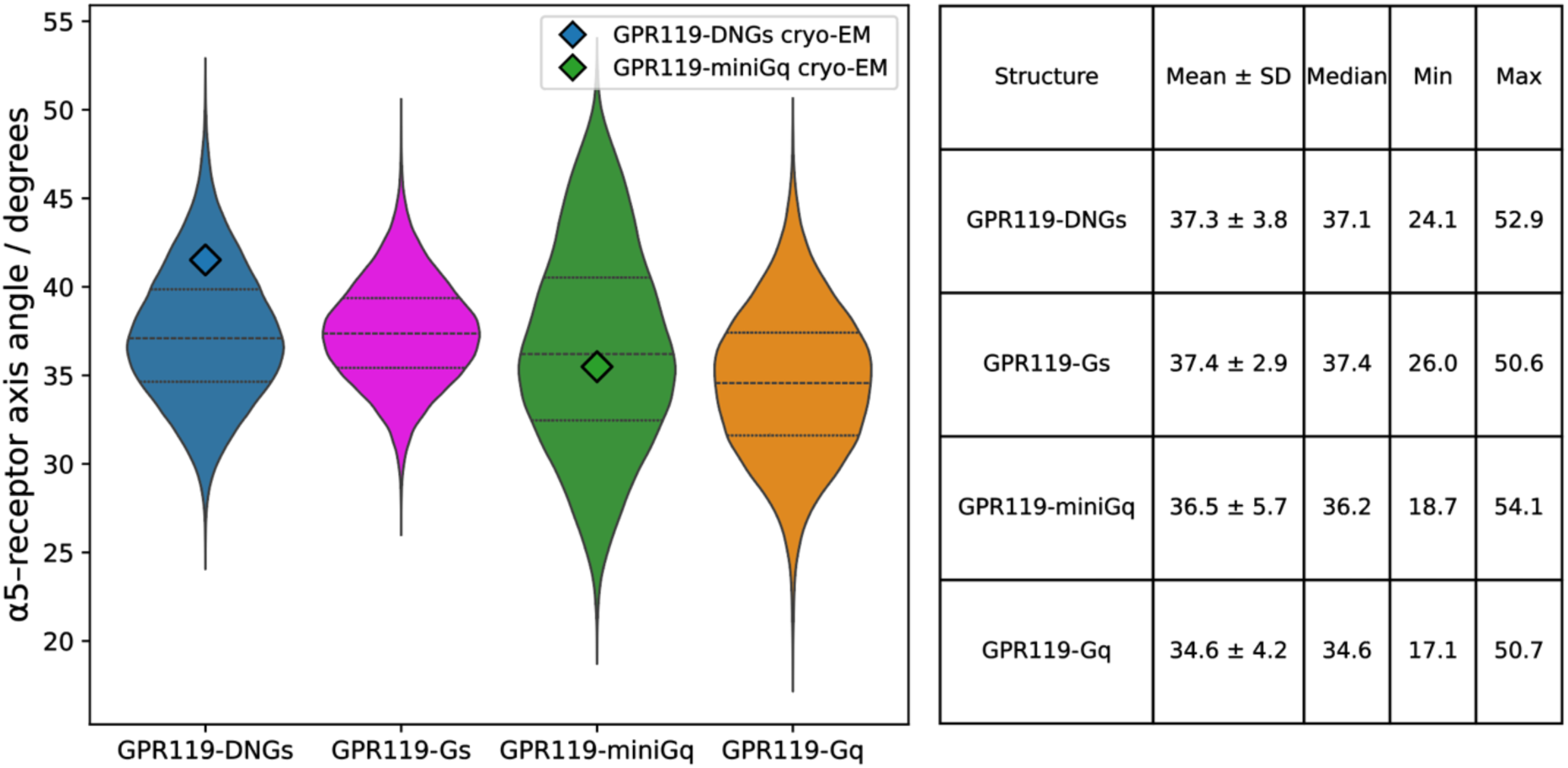
Distributions of the αH5 tilt angle among all the structures across all MD simulations. The table shows the summary of statistics across all structures. Tilt angles from cryo-EM structures are shown as diamonds (GPR119-DNG_s_: 41.5 degrees, GPR119-miniG_q_: 35.5 degrees).

**Supplementary Figure S12.**
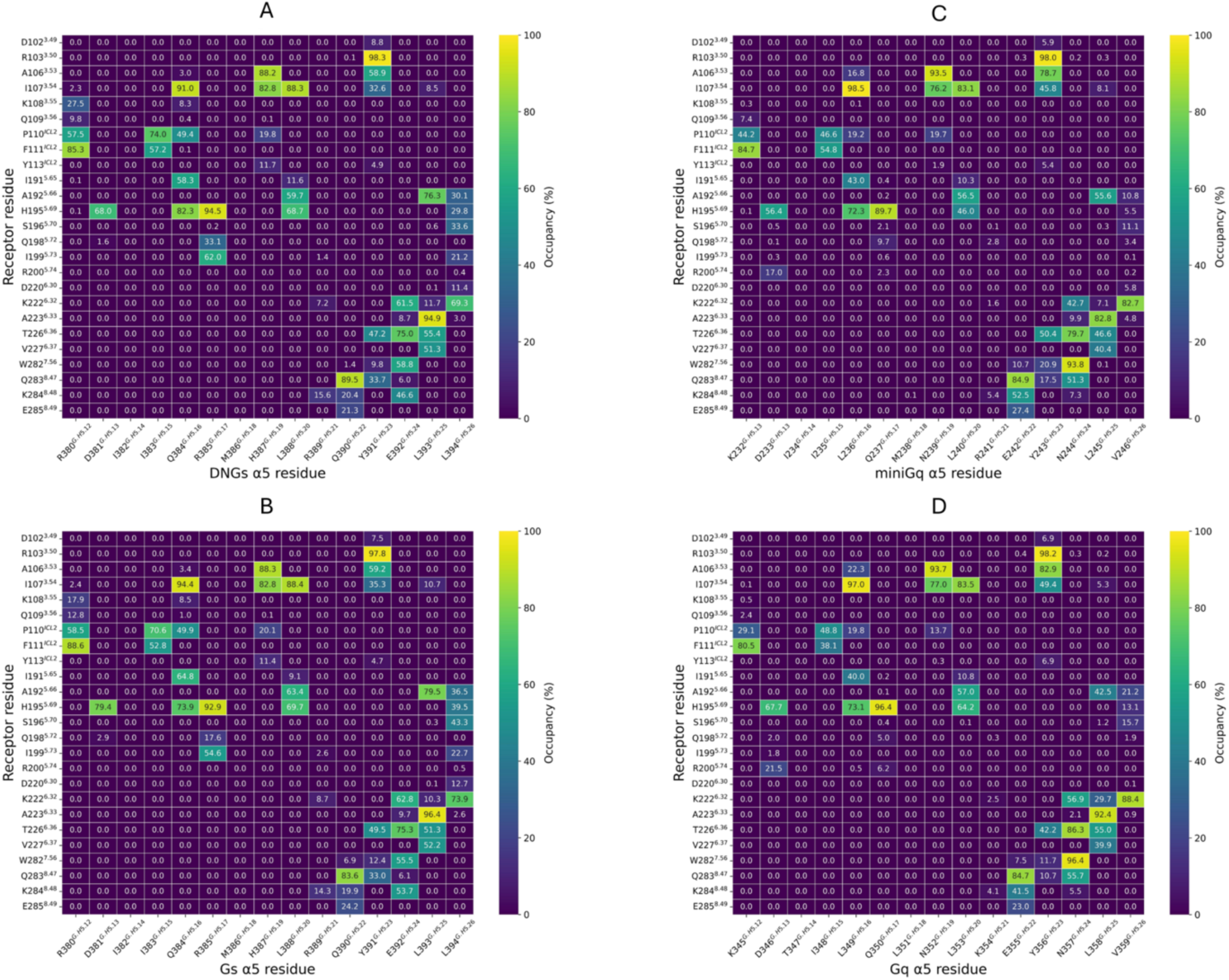
Contact occupancies of direct interactions between GPR119 and the Gα α5 helix, averaged across all MD simulations. (**A**) GPR119-DNG_s_. (**B**) GPR119-G_s_. (**C**) GPR119-miniG_q_. (**D**) GPR119-G_q_. Contact occupancy is defined as the percentage of simulation frames where the closest heavy-atom distance between GPR119 and the α5 helix is ≤ 4 Å.

**Supplementary Figure S13.**
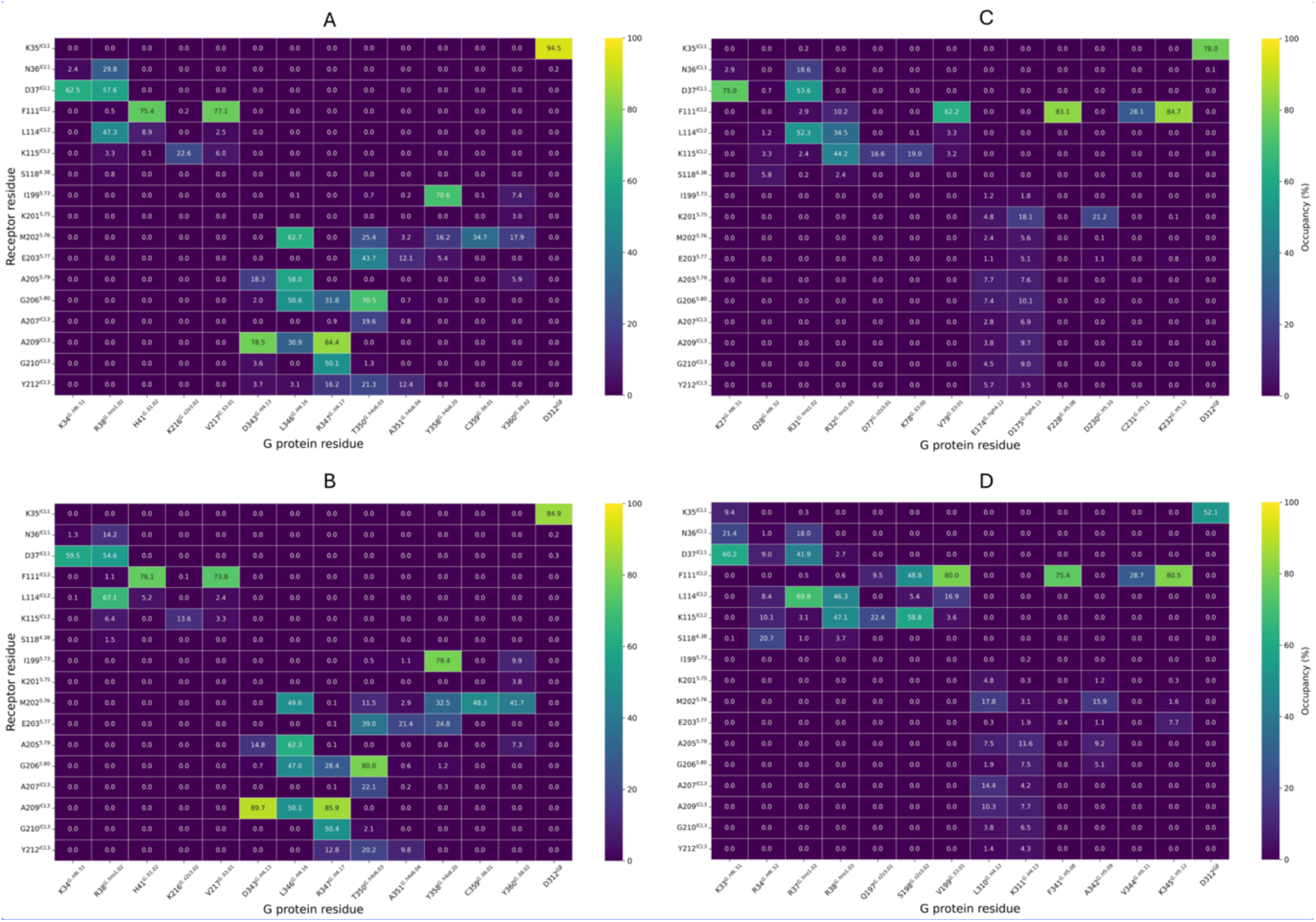
Contact occupancies of direct interactions between GPR119 and G protein residues, excluding the Gα α5 helix, averaged across all MD simulations. (**A**) GPR119-DNG_s_, (**B**) GPR119-G_s_, (**C**) GPR119-miniG_q_, (**D**) GPR119-G_q_. Contact occupancy is defined as the percentage of simulation frames where the closest heavy-atom distance between GPR119 and corresponding G protein is ≤ 4 Å.

**Supplementary Fig. S14.**
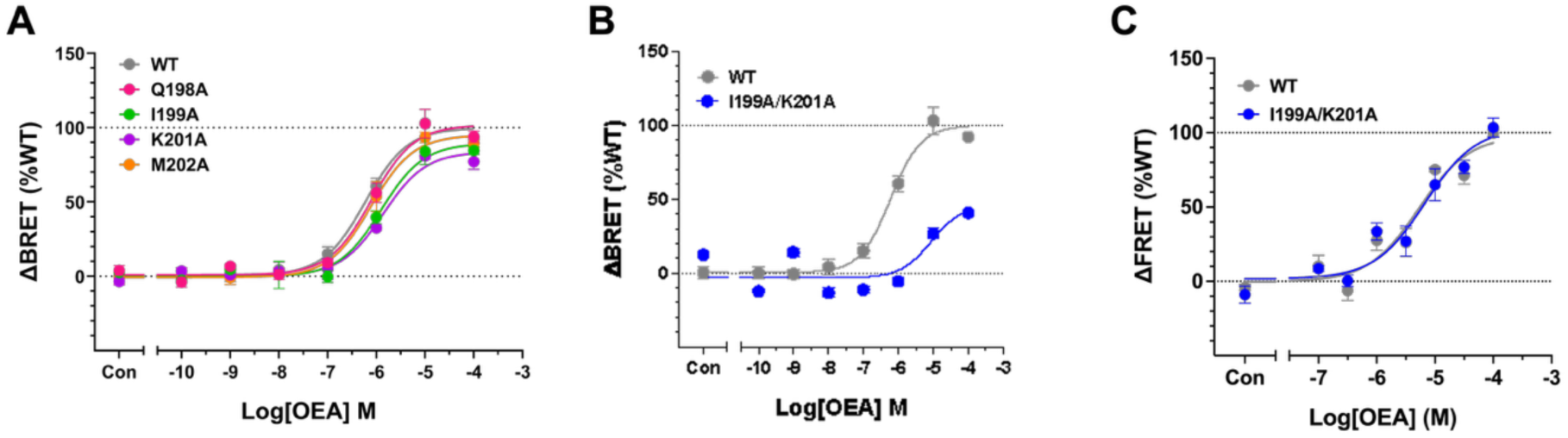
Effects of GPR119 TM5 mutants on G_s_ and G_q_ protein signaling. (**A,B**) Concentration-response curves for OEA-induced G_s_ signaling measured by ONE-GO assay. (**C)** Concentration-response curves for OEA-induced G_q_ signaling assessed by HTRF IP-One assay. Data are normalized to WT and presented as mean ± SEM from three or six (WT) independent experiments. The pEC_50_ and E_max_ values are provided in Supplementary Tables S2 and S3.

**Supplementary Figure S15.**
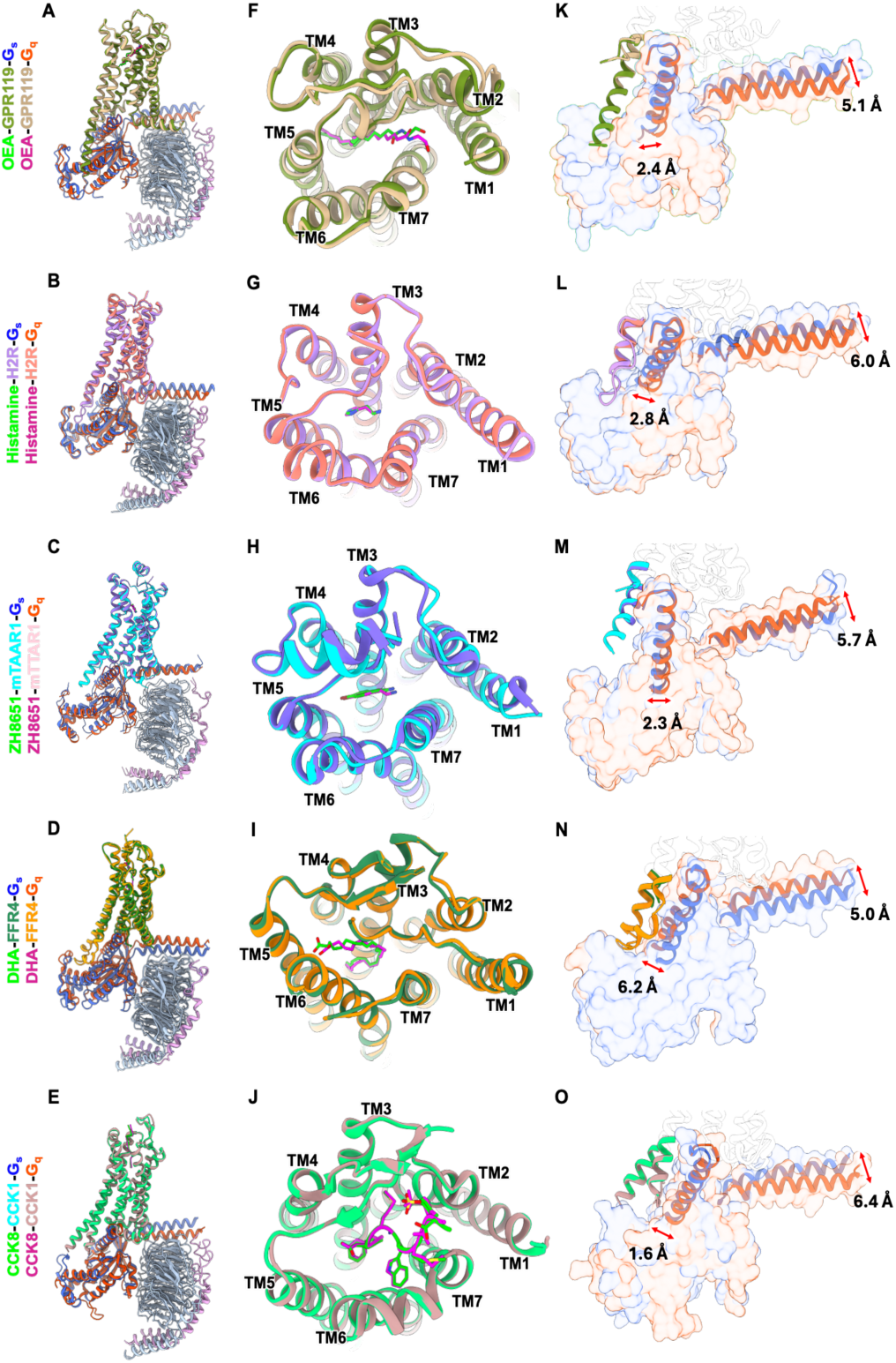
Structural comparison of G_s_- and G_q_-coupled class A GPCRs. (**A**−**E**) Structural superposition of receptor-G protein complexes of GPR119 (A), histamine H2 receptor (H2R, PDB entry 8YN3 and 8YN4) (B), mouse trace amine-associated receptor 1 (mTAAR1, PDB entry 8WC4 and 8WC9) (C), free fatty acid receptor 4 (FFAR4, PDB entry 8H4I and 8H4L) (D), and cholecystokinin A receptor (CCK1R, PDB entry 7EZK and 7EZM) (E) in the G_s_- and G_q_-coupled states. Receptors are shown as cartoons and colored according to the corresponding structures. (**F**−**J**) Extracellular views of the receptor transmembrane bundles highlighting the difference of ligand binding mode between the G_s_- and G_q_-coupled conformations. Bound ligands are shown as sticks. (**K**−**O**) Comparison of TM5−ICL3−TM6 regions and G protein engagement between G_s_ and G_q_-coupled structures. Distances between the corresponding regions in the G_s_- and G_q_-coupled structures are indicated. Structures were aligned based on the Cα atoms of the receptor transmembrane helices.

**Supplementary Figure S16.**
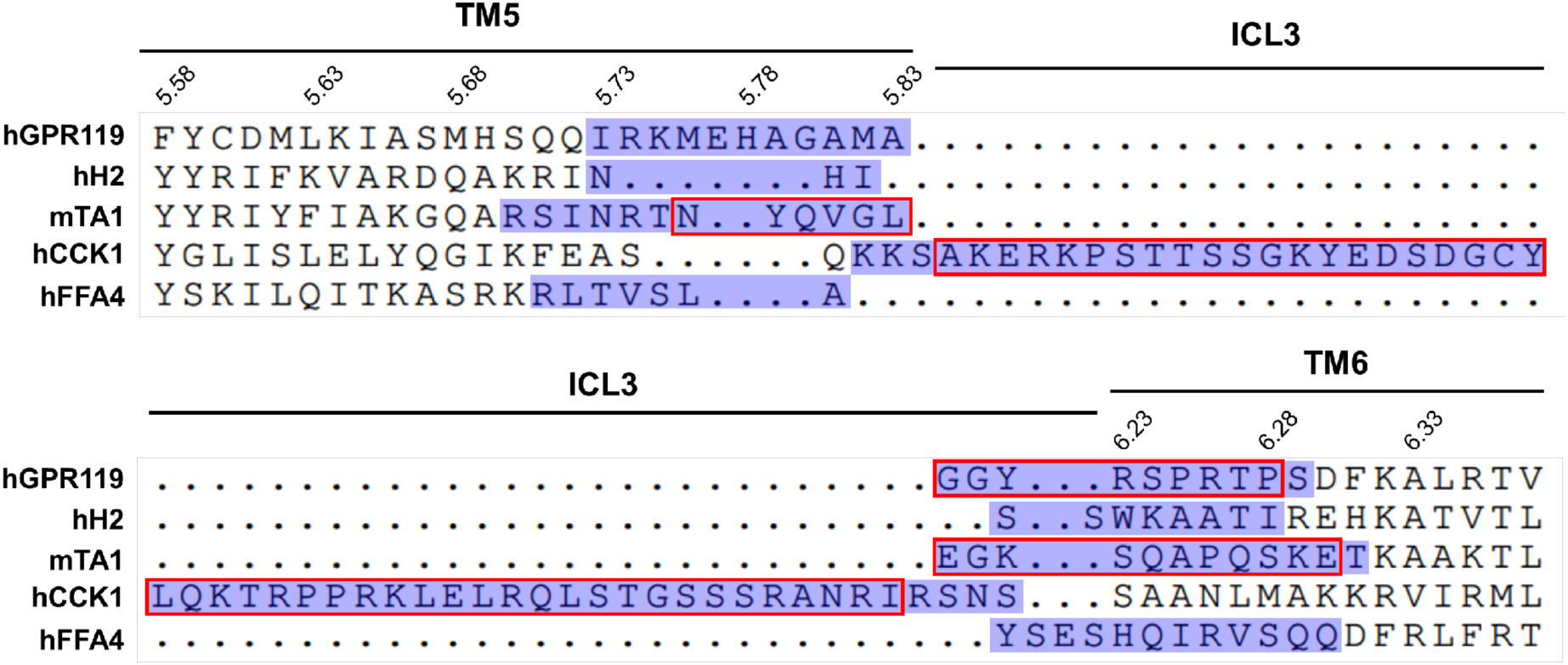
Sequence alignment of the TM5–ICL3–TM6 regions among representative class A GPCRs. The alignment highlights the disordered TM5–ICL3–TM6 regions of receptors complexed with G_s_ or G_q_ proteins. The disordered regions in the primary G protein-coupled structures (hGPR119-G_s_, hH2-G_s_, mTA1-G_s_, hFFA4-G_q_, and hCCK1-G_q_) are outlined in red boxes, whereas the disordered regions in the secondary G protein-coupled structures (hGPR119-G_q_, hH2-G_q_, mTA1-G_q_, hFFA4-G_s_, and hCCK1-G_s_) are highlighted in blue, indicating more extensive disordered regions.

**Supplementary Figure S17.**
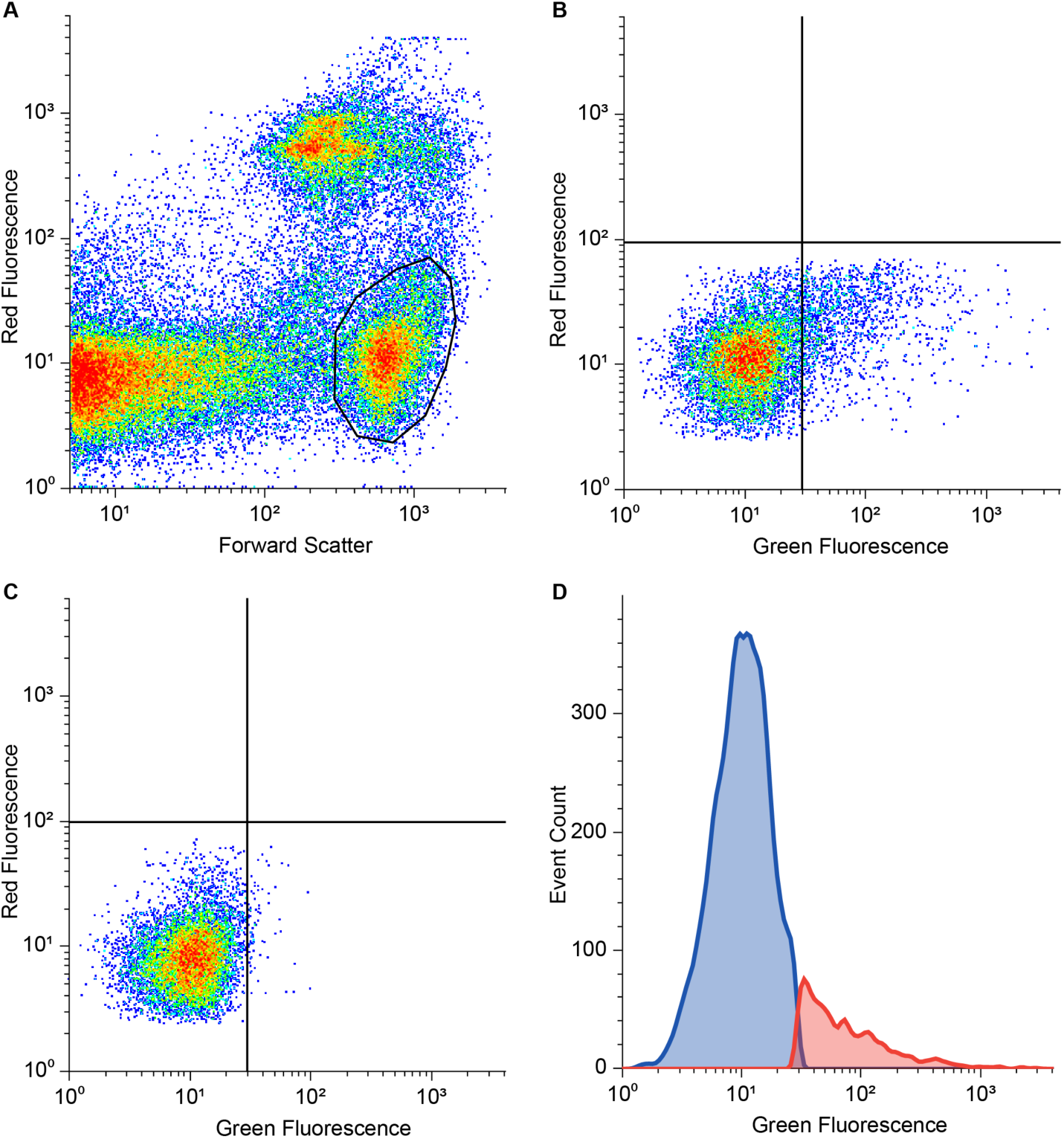
Gating strategy used for measuring GPR119 surface expression by flow cytometry. (**A**) Live cells were selected on the plot of Red Fluorescence (7-AAD) vs Forward Scatter. (**B, C**) Green fluorescence (Anti-FLAG M2-FITC) threshold was selected using cells expressing WT GPR119 (positive control, B) and the same cells transfected with empty vector (negative control, C). (**D**) Histogram of events recorded in the Green Fluorescence channel for WT GPR119. The population of cells expressing GPR119 is colored red.

**Supplementary Table S1.** Cryo-EM data collection, refinement, and validation statistics.

|  | **OEA-GPR119-G<sub>s</sub>-Nb35** | **OEA-GPR119-miniG<sub>q</sub>** |
| --- | --- | --- |
| **PDB entry** | 36QE | 36QF |
| **EMDB entry** |  |  |
| Composite map | EMD-77772 | EMD-77773 |
| Consensus map | EMD-77768 | EMD-77770 |
| Focused map (Receptor) | EMD-77769 | EMD-77771 |
| **Data collection and processing** |  |  |
| Magnification | 105,000× | 105,000× |
| Voltage (kV) | 300 | 300 |
| Electron exposure (e<sup>-</sup>/Å<sup>2</sup>) | 61 | 60 |
| Defocus range (μm) | -0.8 to -2.2 | -0.8 to -2.2 |
| Pixel size (Å) | 0.85 | 0.85 |
| Symmetry imposed | C1 | C1 |
| Initial particle images (no.) | 710,476 | 2,913,751 |
| Final particle images (no.) | 337,012 | 557,856 |
| Map resolution (Å) | 2.72 | 3.05 |
| FSC threshold | 0.143 | 0.143 |
| Map resolution range (Å) | 2.0 – 5.0 | 2.0 – 5.0 |
| **Refinement** |  |  |
| Initial model used (PDB code) | 8VHF | 8VHF |
| Model resolution (Å) | 2.8 | 3.2 |
| FSC threshold | 0.5 | 0.5 |
| Map sharpening *B* factor (Å<sup>2</sup>) | -84.2 | -109.5 |
| **Model composition** |  |  |
| Non-hydrogen atoms | 8,245 | 7,162 |
| Protein residues | 1,047 | 917 |
| Ligands | 2 | 1 |
| Water | 9 | 2 |
| ***B* factors (Å<sup>2</sup>)** |  |  |
| Protein | 65.1 | 69.2 |
| Ligand | 79.2 | 44.9 |
| Water | 56.1 | 45.3 |
| **R.m.s. deviations** |  |  |
| Bond lengths (Å) | 0.002 | 0.002 |
| Bond angles (°) | 0.518 | 0.456 |
| **Validation** |  |  |
| MolProbity score | 1.03 | 1.15 |
| Clashscore | 2.44 | 3.57 |
| Poor rotamers (%) | 0.57 | 0.13 |
| **Ramachandran plot** |  |  |
| Favored (%) | 98.16 | 98.56 |
| Allowed (%) | 1.84 | 1.44 |
| Disallowed (%) | 0.0 | 0.0 |

**Supplementary Table S2.**
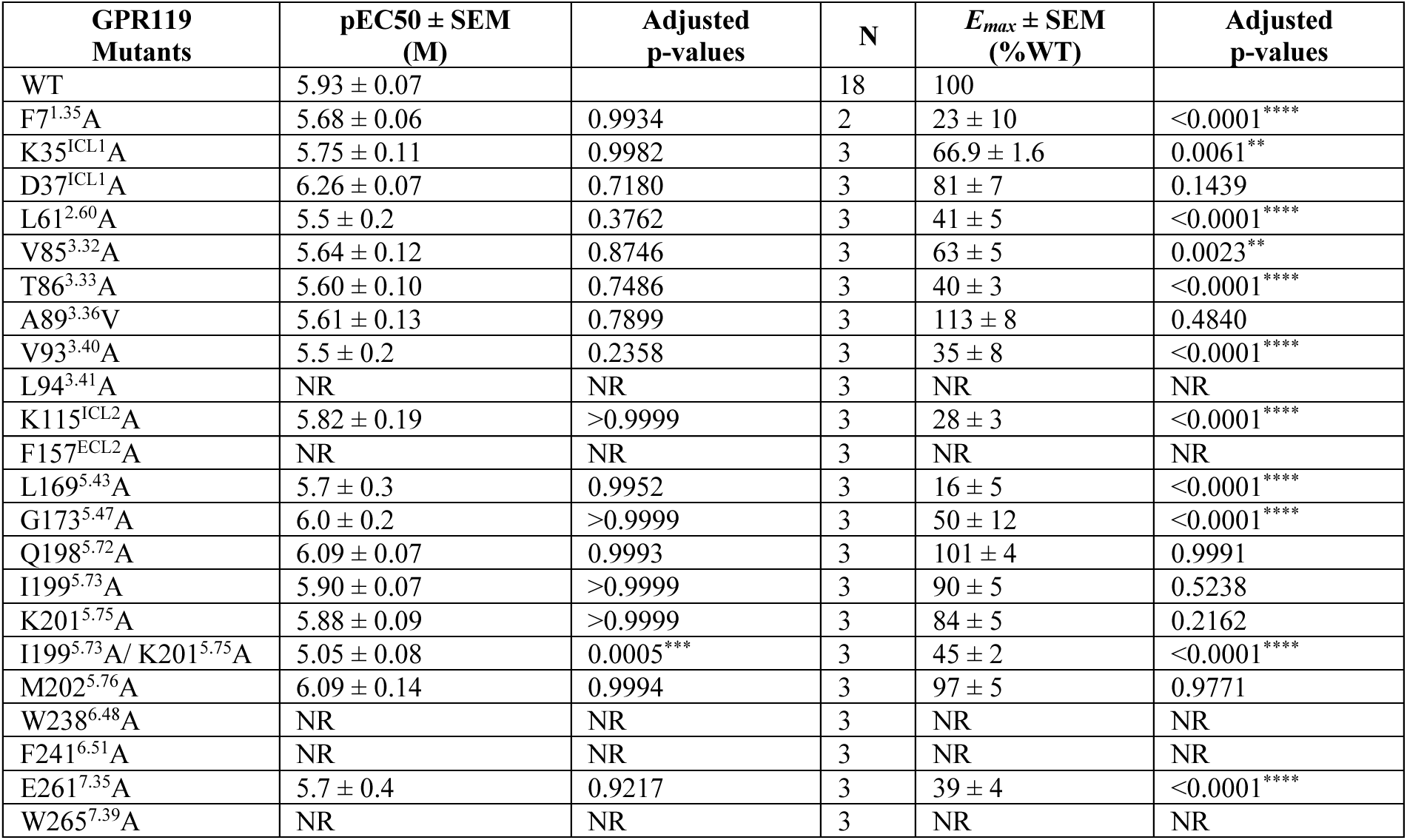
Effect of GPR119 mutants on OEA-induced G_s_ signaling. G_s_ activation was measured by the ONE-GO G_s_ coupling assay. The mutant signal was normalized to WT (set to 100%). p*EC*_50_ and *E_max_* values represent mean ± SEM from N independent experiments conducted in triplicates. NR indicates no response. Significant difference relative to WT was determined by one-way ANOVA with Dunnett’s multiple comparisons test as p < 0.05 (*), p < 0.001 (**), and p < 0.0001 (****).

**Supplementary Table S3.**
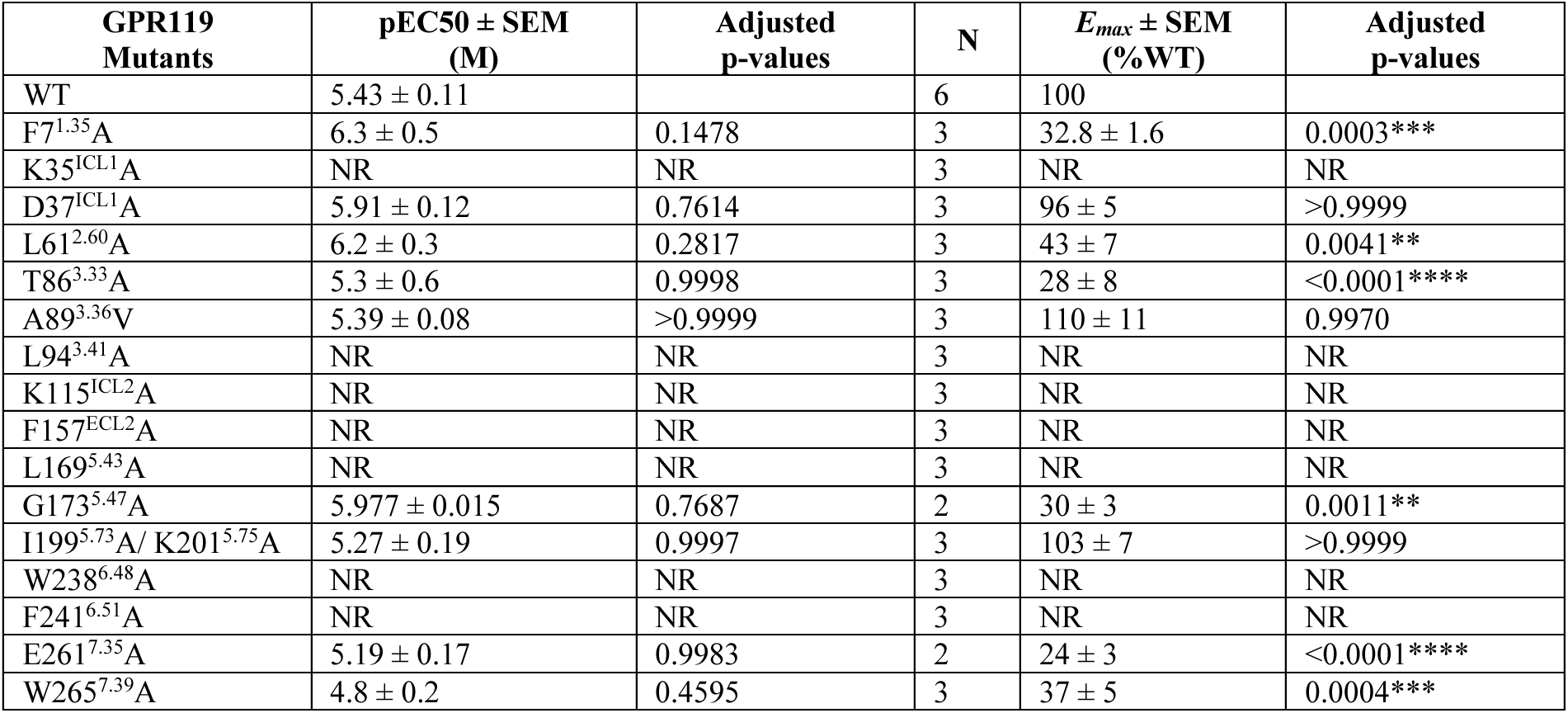
Effect of GPR119 mutants on OEA-induced G_q_ signaling. G_q_ activation was measured by the HTRF IP-ONE assay. The mutant signal was normalized to WT (set to 100%). p*EC*_50_ and *E_max_* values represent mean ± SEM from N independent experiments conducted in triplicates. NR indicates no response. Significant difference relative to WT was determined by one-way ANOVA with Dunnett’s multiple comparisons test. Significant p-values are labeled as ** (p < 0.001), *** (p < 0.001), and **** (p < 0.0001).

**Supplementary Table S4.** Cell surface expression of WT GPR119 and mutants, measured by flow cytometry. The median signal of mutant-expressing live cells is normalized to that of the WT receptor (set to 100%). The surface expression values represent mean ± SEM from three independent experiments. Significant differences relative to WT were determined by mixed effects one-way ANOVA with Dunnett’s multiple comparisons test conducted before normalization and using a log-normal distribution.

| GPR119 Mutants | Surface expression, %WT mean ± SEM | Adjusted p-values |
| --- | --- | --- |
| WT | 100 | N/A |
| F7<sup>1.35</sup>A | 95 ± 6 | 0.9761 |
| K35<sup>ICL1</sup>A | 94 ± 5 | 0.8814 |
| D37<sup>ICL1</sup>A | 96 ± 4 | 0.8960 |
| L61<sup>2.60</sup>A | 90.9 ± 1.4 | 0.0948 |
| V85<sup>3.32</sup>A | 91 ± 8 | 0.9110 |
| T86<sup>3.33</sup>A | 87 ± 4 | 0.3877 |
| A89<sup>3.36</sup>V | 99 ± 9 | >0.9999 |
| V93<sup>3.40</sup>A | 95 ± 8 | 0.9928 |
| L94<sup>3.41</sup>A | 90 ± 7 | 0.7788 |
| K115<sup>ICL2</sup>A | 88 ± 6 | 0.5661 |
| F157<sup>ECL2</sup>A | 85 ± 6 | 0.4639 |
| L169<sup>5.43</sup>A | 92 ± 4 | 0.5661 |
| G173<sup>5.47</sup>A | 94 ± 4 | 0.8401 |
| Q198<sup>5.72</sup>A | 93 ± 7 | 0.8800 |
| I199<sup>5.73</sup>A | 98 ± 5 | 0.9999 |
| K201<sup>5.75</sup>A | 92 ± 5 | 0.7125 |
| I199<sup>5.73</sup>A/ K201<sup>5.75</sup>A | 97 ± 5 | 0.9977 |
| M202<sup>5.76</sup>A | 94 ± 5 | 0.8119 |
| W238<sup>6.48</sup>A | 91 ± 6 | 0.7871 |
| F241<sup>6.51</sup>A | 95.0 ± 1.4 | 0.2572 |
| E261<sup>7.35</sup>A | 93 ± 5 | 0.7272 |
| W265<sup>7.39</sup>A | 88 ± 6 | 0.5873 |

## References

1. S. Padhi, A. K. Nayak, A. Behera, Type II diabetes mellitus: a review on recent drug based therapeutics. Biomed Pharmacother 131, 110708 (2020).

2. R. E. Ramírez-Orozco et al., Potential metabolic and behavioural roles of the putative endocannabinoid receptors GPR18, GPR55 and GPR119 in feeding. Current neuropharmacology 17, 947–960 (2019).

3. H. Overton, M. Fyfe, C. Reynet, GPR119, a novel G protein-coupled receptor target for the treatment of type 2 diabetes and obesity. British journal of pharmacology 153, S76–S81 (2008).

4. Z.-L. Chu et al., A role for intestinal endocrine cell-expressed g protein-coupled receptor 119 in glycemic control by enhancing glucagon-like Peptide-1 and glucose-dependent insulinotropic Peptide release. Endocrinology 149, 2038–2047 (2008).

5. H. Liu, A. Dear, L. Knudsen, R. Simpson, A long-acting GLP-1 analogue attenuates induction of PAI-1 and vascular adhesion molecules. J Endocrinol 201, 59–66 (2009).

6. H. S. Hansen, M. M. Rosenkilde, J. J. Holst, T. W. Schwartz, GPR119 as a fat sensor. Trends in pharmacological sciences 33, 374–381 (2012).

7. T. Ohishi, S. Yoshida, The therapeutic potential of GPR119 agonists for type 2 diabetes. Expert opinion on investigational drugs 21, 321–328 (2012).

8. D.-S. Im, GPR119 and GPR55 as receptors for fatty acid ethanolamides, oleoylethanolamide and palmitoylethanolamide. International journal of molecular sciences 22, 1034 (2021).

9. H. S. Hansen, T. A. Diep, N-acylethanolamines, anandamide and food intake. Biochemical pharmacology 78, 553–560 (2009).

10. M. Igarashi et al., Intestinal GPR119 activation by microbiota-derived metabolites impacts feeding behavior and energy metabolism. Molecular Metabolism 67, 101649 (2023).

11. H. A. Hassing et al., Biased signaling of lipids and allosteric actions of synthetic molecules for GPR119. Biochemical pharmacology 119, 66–75 (2016).

12. M. Zhang et al., G protein-coupled receptors (GPCRs): advances in structures, mechanisms and drug discovery. Signal transduction and targeted therapy 9, 88 (2024).

13. M. Hauge et al., Gq and Gs signaling acting in synergy to control GLP-1 secretion. Molecular and Cellular Endocrinology 449, 64–73 (2017).

14. Y. Qian et al., Activation and signaling mechanism revealed by GPR119-Gs complex structures. Nature communications 13, 7033 (2022).

15. P. Xu et al., Structural identification of lysophosphatidylcholines as activating ligands for orphan receptor GPR119. Nature structural & molecular biology 29, 863–870 (2022).

16. R. Li et al., Structure of human GPR119-Gs complex binding APD597 and characterization of GPR119 binding agonists. Frontiers in Pharmacology 15, 1310231 (2024).

17. E. Chun et al., Fusion partner toolchest for the stabilization and crystallization of G protein-coupled receptors. Structure 20, 967–976 (2012).

18. J. Duan et al., Cryo-EM structure of an activated VIP1 receptor-G protein complex revealed by a NanoBiT tethering strategy. Nat Commun 11, 4121 (2020).

19. Y.-L. Liang et al., Phase-plate cryo-EM structure of a biased agonist-bound human GLP-1 receptor–Gs complex. Nature 555, 121–125 (2018).

20. X. Li et al., Structural basis for ligand recognition and activation of the prostanoid receptors. Cell Reports 43, (2024).

21. S. Maeda et al., Development of an antibody fragment that stabilizes GPCR/G-protein complexes. Nature communications 9, 3712 (2018).

22. P. Laleh, K. Yaser, O. Alireza, Oleoylethanolamide: A novel pharmaceutical agent in the management of obesity-an updated review. Journal of Cellular Physiology 234, 7893–7902 (2019).

23. K. Proulx et al., Mechanisms of oleoylethanolamide-induced changes in feeding behavior and motor activity. American Journal of Physiology-Regulatory, Integrative and Comparative Physiology 289, R729–R737 (2005).

24. L. M. Lauffer, R. Iakoubov, P. L. Brubaker, GPR119 is essential for oleoylethanolamide-induced glucagon-like peptide-1 secretion from the intestinal enteroendocrine L-cell. Diabetes 58, 1058–1066 (2009).

25. Z. Ouyang et al., Structural insights into the activation of G protein-coupled receptor 119 by the agonist GSK1292263 and ligands selectivity among novel cannabinoid receptors. International Journal of Biological Macromolecules 319, 145476 (2025).

26. T. Flock et al., Universal allosteric mechanism for Gα activation by GPCRs. Nature 524, 173–179 (2015).

27. X. Zhang et al., Structural basis of ligand recognition and activation of the histamine receptor family. Nature Communications 15, 8296 (2024).

28. P. Shang et al., Structural and signaling mechanisms of TAAR1 enabled preferential agonist design. Cell 186, 5347–5362. e5324 (2023).

29. H. Yin et al., Structural basis of omega-3 fatty acid receptor FFAR4 activation and G protein coupling selectivity. Cell Research 33, 644–647 (2023).

30. Q. Liu et al., Ligand recognition and G-protein coupling selectivity of cholecystokinin A receptor. Nature chemical biology 17, 1238–1244 (2021).

31. D. J. Drucker, Mechanisms of action and therapeutic application of glucagon-like peptide-1. Cell metabolism 27, 740–756 (2018).

32. J. Ghislain, V. Poitout, Targeting lipid GPCRs to treat type 2 diabetes mellitus—Progress and challenges. Nature Reviews Endocrinology 17, 162–175 (2021).

33. D. Mears, Regulation of insulin secretion in islets of Langerhans by Ca2+ channels. The Journal of membrane biology 200, 57–66 (2004).

34. D. Billups, B. Billups, R. J. Challiss, S. R. Nahorski, Modulation of Gq-protein-coupled inositol trisphosphate and Ca2+ signaling by the membrane potential. Journal of Neuroscience 26, 9983–9995 (2006).

35. Z. Rankovic, T. F. Brust, L. M. Bohn, Biased agonism: An emerging paradigm in GPCR drug discovery. Bioorganic & medicinal chemistry letters 26, 241–250 (2016).

36. O. S. Oduori et al., Gs/Gq signaling switch in β cells defines incretin effectiveness in diabetes. The Journal of clinical investigation 130, 6639–6655 (2020).

37. M. Crutchlow et al., Combined GPR40 and GPR119 full agonism with K-757 and K-833 results in robust glucose lowering and modest weight loss in overweight/obese subjects with T2DM. Diabetes, Obesity and Metabolism 27, 6264–6274 (2025).

38. D. Kim, W. Liu, R. Viner, V. Cherezov, Native mass spectrometry prescreening of G protein-coupled receptor complexes for cryo-EM structure determination. Structure 32, 2206–2219. e2204 (2024).

39. Y. L. Liang et al., Phase-plate cryo-EM structure of a biased agonist-bound human GLP-1 receptor-Gs complex. Nature 555, 121–125 (2018).

40. R. Nehme et al., Mini-G proteins: Novel tools for studying GPCRs in their active conformation. PloS one 12, e0175642 (2017).

41. P. Liu et al., The structural basis of the dominant negative phenotype of the Gαi1β1γ2 G203A/A326S heterotrimer. Acta Pharmacologica Sinica 37, 1259–1272 (2016).

42. B. Davoudinasab et al., Structural insights into the mechanism of activation and inhibition of the prostaglandin D2 receptor 1. Nature Communications 16, 8944 (2025).

43. Z. Zhang, Y. Wang, Y. Ding, M. Hattori, Structure-based engineering of anti-GFP nanobody tandems as ultra-high-affinity reagents for purification. Sci Rep 10, 6239 (2020).

44. A. Punjani, J. L. Rubinstein, D. J. Fleet, M. A. Brubaker, cryoSPARC: algorithms for rapid unsupervised cryo-EM structure determination. Nat Methods 14, 290–296 (2017).

45. D. Liebschner et al., Macromolecular structure determination using X-rays, neutrons and electrons: recent developments in Phenix. Biological Crystallography 75, 861–877 (2019).

46. A. Punjani, J. L. Rubinstein, D. J. Fleet, M. A. Brubaker, cryoSPARC: algorithms for rapid unsupervised cryo-EM structure determination. Nature methods 14, 290–296 (2017).

47. E. F. Pettersen et al., UCSF Chimera--a visualization system for exploratory research and analysis. J Comput Chem 25, 1605–1612 (2004).

48. P. D. Adams et al., PHENIX: a comprehensive Python-based system for macromolecular structure solution. Acta Crystallogr D Biol Crystallogr 66, 213–221 (2010).

49. P. Emsley, K. Cowtan, Coot: model-building tools for molecular graphics. Acta Crystallogr D Biol Crystallogr 60, 2126–2132 (2004).

50. A. Raskovalov, D. Kim, V. Cherezov, ONE-GO: Direct detection of context-dependent GPCR activity. Cell Research 34, 543–544 (2024).

51. R. Janicot et al., Direct interrogation of context-dependent GPCR activity with a universal biosensor platform. Cell 187, 1527–1546. e1525 (2024).

52. M. Abraham. et al., GROMACS 2026.0 Source code (2026).

53. J. Huang, A. D. MacKerell Jr, CHARMM36 all-atom additive protein force field: Validation based on comparison to NMR data. Journal of computational chemistry 34, 2135–2145 (2013).

54. S. Jo, T. Kim, V. G. Iyer, W. Im, CHARMM-GUI: a web-based graphical user interface for CHARMM. Journal of computational chemistry 29, 1859–1865 (2008).

55. B. R. Brooks et al., CHARMM: the biomolecular simulation program. Journal of computational chemistry 30, 1545–1614 (2009).

56. J. Lee et al., CHARMM-GUI input generator for NAMD, GROMACS, AMBER, OpenMM, and CHARMM/OpenMM simulations using the CHARMM36 additive force field. Biophysical journal 110, 641a (2016).

57. M. Sandhu et al., Conformational plasticity of the intracellular cavity of GPCR− G-protein complexes leads to G-protein promiscuity and selectivity. Proceedings of the National Academy of Sciences 116, 11956–11965 (2019).

58. D. Wang, H. Yu, X. Liu, J. Liu, C. Song, The orientation and stability of the GPCR-Arrestin complex in a lipid bilayer. Scientific reports 7, 16985 (2017).

59. R. T. McGibbon et al., MDTraj: a modern open library for the analysis of molecular dynamics trajectories. Biophysical journal 109, 1528–1532 (2015).

60. G. Pérez-Hernández, P. W. Hildebrand, Mdciao: accessible analysis and visualization of molecular dynamics simulation data. PLoS computational biology 21, e1012837 (2025).

